# Connexin 43 confers chemoresistance through activating PI3K

**DOI:** 10.1101/2020.10.14.339275

**Authors:** Kevin J Pridham, Farah Shah, Kevin L Sheng, Sujuan Guo, Min Liu, Pratik Kanabur, Samy Lamouille, Gabrielle Lewis, Marc Morales, Jane Jourdan, Christina L Grek, Gautam G Ghatnekar, Robin Varghese, Deborah F Kelly, Robert G Gourdie, Zhi Sheng

**Affiliations:** Fralin Biomedical Research Institute at VTC, Roanoke, VA 24016, USA; Department of Biomedical Affairs and Research, Edward Via College of Osteopathic Medicine, Blacksburg, VA 24060, USA; Department of Internal Medicine, Virginia Tech Carilion School of Medicine, Roanoke, VA 24016, USA; Department of Basic Science Education, Virginia Tech Carilion School of Medicine, Roanoke, VA 24016, USA; Department of Biological Sciences, Virginia Tech, Blacksburg VA 24061, USA; FirstString Research, Inc., Mount Pleasant, SC 29464, USA; Department of Biomedical Engineering, Pennsylvania State University, University Park, PA 16802, USA; Huck Institutes of the Life Sciences, Pennsylvania State University, University Park, PA 16802, USA; Center for Structural Oncology, Pennsylvania State University, University Park, PA 16802, USA; Department of Emergency Medicine, Virginia Tech Carilion School of Medicine, Roanoke, VA 24016, USA; Faculty of Health Science, Virginia Tech, Blacksburg, VA 24061, United States

**Keywords:** Cx43, PI3K, chemoresistance, PIK3CB/p110β, temozolomide, glioblastoma

## Abstract

Circumventing chemoresistance is crucial for effectively treating glioblastoma due to limited therapeutic options. The gap junction protein connexin 43 (Cx43) renders glioblastoma resistant to the frontline chemotherapy temozolomide; however, targeting Cx43 is difficult because mechanisms underlying Cx43-mediated chemoresistance remain elusive. Here we show that Cx43, but not other connexins, is highly expressed in glioblastoma and strongly correlates with poor patient prognosis and chemoresistance, making Cx43 the prime therapeutic target among all connexins. The intracellular carboxyl terminus of Cx43 binds to phosphatidylinositol 3-kinase (PI3K) catalytic subunit β (PIK3CB, also called PI3Kβ or p110β), thereby activating PI3K signaling independent of Cx43-channels and subsequently inducing temozolomide resistance. A combination of αCT1, a Cx43-targeting peptide inhibitor, and PIK3CB-selective inhibitors restores temozolomide sensitivity *in vitro* and *in vivo*. This study not only reveals novel mechanistic insights into chemoresistance in glioblastoma, but also demonstrates that targeting Cx43 and PIK3CB/p110β is an effective approach for overcoming chemoresistance.

---

Overcoming resistance to chemotherapy such as temozolomide (TMZ) has proven perplexing and remains a key unmet clinical need. As an alkylating agent, TMZ reacts with DNA at multiple sites, yielding O^6^-methylguanine lesions that subsequently induce DNA breaks and eventually cell death (1). Given that TMZ is able to pass the blood-brain barrier (2), this drug has been used as the frontline chemotherapy for glioblastoma (GBM), an aggressive and lethal cancer that accounts for approximately half of all malignant brain tumors and has a grim prognosis with an average survival time of 14.6 months (3, 4). Adding to this dismal outcome, nearly 90% of patients with GBM succumb to tumor recurrence and the average survival for recurrent GBM is about 5.5-7.5 months due to limited therapeutic options and resistance to TMZ (5). Hence, overcoming TMZ resistance is key to effectively treating GBM and curbing GBM progression. Poor responses of nearly 50% of GBM patients to TMZ are due to the expression of O-6-methylguanine-DNA methyltransferase (MGMT) (6, 7). MGMT repairs TMZ-induced DNA damage, conferring MGMT-dependent TMZ resistance; as such, inhibiting MGMT has shown encouraging clinical benefits (8). Patients with no MGMT expression also develop MGMT-independent resistance to TMZ (9, 10). Factors involved in MGMT-independent TMZ resistance include the DNA mismatch repair pathway and genetic alterations (11, 12). However, targeting these factors to circumvent TMZ resistance has been a daunting task. Deeper insights into MGMT-independent TMZ resistance are therefore needed.

Recently, several lines of evidence have indicated that the gap junction protein connexin 43 (Cx43; also known as gap junction protein A1, *GJA1*), a channel-forming protein important for intercellular communication (13), controls the response of GBM cells to TMZ. Ectopic expression of Cx43 renders GBM cells resistant to TMZ (14–17) and blocking Cx43 using different approaches such as antibodies or channel inhibitors restores TMZ sensitivity (14–20). However, it remains unclear whether Cx43-mediated TMZ resistance depends on MGMT. Our recent work (21) reveals that high levels of Cx43 in MGMT-deficient GBM cell lines and primary patient samples correlate with poor responses to TMZ and that αCT1, a clinically-tested therapeutic peptide that comprises the Cx43 carboxyl terminus (CT) and an antennapedia cell-penetrating sequence (22), antagonizes TMZ resistance. Our results have been verified by an independent study using nanoparticle-conjugated αCT1 (23). Nonetheless, the molecular underpinnings of Cx43-mediated TMZ resistance remains elusive, making it difficult to effectively target Cx43 to treat GBM.

In this report, we determined the role of connexins in GBM prognosis and TMZ resistance, explored how Cx43 activates phosphatidylinositol-3 kinase (PI3K) independent of Cx43 channels and induces TMZ resistance, and examined a candidate triple combinational therapy entailing the Cx43 inhibitor αCT1, PI3K-selective inhibitors, and TMZ in preclinical studies for its effectiveness in overcoming TMZ resistance.

## Results

### Cx43, but not other connexins, is highly expressed in GBM and correlates with poor prognosis and chemoresistance

There are 21 known connexins (Supplemental Table 1). Whether all these connexins are equally important for GBM survival and chemoresistance has not yet been explored. To address this, we queried publicly available online GBM databases including: The Cancer Genome Atlas (TCGA; https://www.cancer.gov/tcga), GlioVis (24), Chinese Glioma Gene Atlas (CGGA), and the Cancer Dependency Map (DepMap) (25). Cx43 mRNA was consistently expressed at the highest level among all connexins in primary GBM tumors from six different datasets (**Fig. 1A-E** and Supplemental Fig. S1A-C) and 54 GBM cell lines (Supplemental Fig. S1D). Notable, despite that different connexins were detected in these studies, levels of Cx43 mRNA were significantly higher than other connexins (*P* < 0.0001). Based on immunostaining results retrieved from The Human Protein Atlas (26), levels of Cx43 protein in high-grade glioma were also significantly higher than other connexins, except Cx37 or Cx40 (**Fig. 1F**). In Cx43-high tumors, other connexins were scored as either not detected, low, or medium in the same tumor (**Fig. 1G** and Supplemental Fig. S2), suggestive of a dominant expression of Cx43. Collectively, Cx43 is expressed at the highest level among all connexins in GBM and high-grade glioma.

**Fig. 1.**
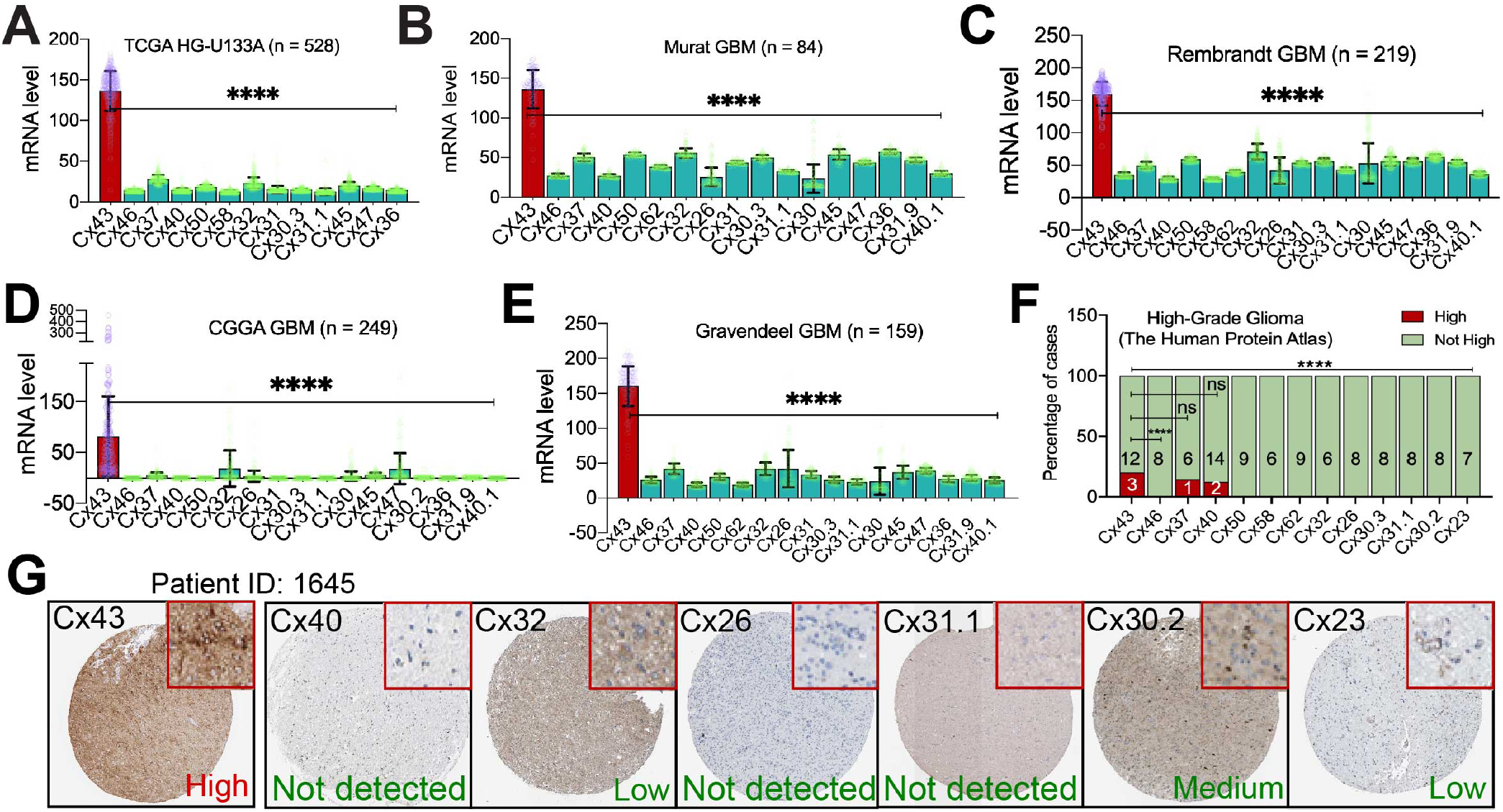
Cx43 is expressed at the highest level among all connexins in GBM. mRNA levels of connexins in GBMs from The Cancer Genome Atlas (TCGA; **A**), Murat (**B**), Rembrandt (**C**), Chinese Glioma Gene Altas (CGGA; **C**), and Gravendeel (**E**). Shown are average reads of microarray or RNAseq. Cx43 is presented as red bars with purple data points. Other connexins are labeled as green bars and yellow data points. Error bars are either standard deviations or standard errors. (**F**) Staining scores of connexins in high-grade glioma. Case numbers with high (red) or not high (green) levels of connexins are shown. (**G**) Histological images of connexins in a high-grade glioma tumor. Inset images (highlighted in red) were cropped from original images in order to highlight immunostaining details. GBM datasets were retrieved from cBioPortal, GlioVis, or CGGA data portal. Immunostaining results of high-grade glioma were obtained from the Human Protein Atlas. Statistical analyses: One-Way ANOVA and Fisher’s exact test. ns: not significant; ****: *P* < 0.0001.

Kaplan-Meier analyses (**Fig. 2A** and Supplemental Fig. S3A) revealed that high levels of Cx43 mRNA were associated with poor prognosis of GBM patients (All GBM). However, the lifespan of Cx43-high primary GBM was not significantly shorter than Cx43-low primary GBM (Primary GBM, *P* > 0.05). This is perhaps due to the fact that 50% of primary GBM express MGMT (6) and that Cx43 correlates with the survival of MGMT-deficient patients (21). Indeed, in MGMT-deficient/TMZ-untreated primary GBMs, high levels of Cx43 correlated with poor prognosis (MGMT–/TMZ–, *P* < 0.05), whereas Cx43 levels had no relationship with the survival of MGMT+/TMZ– GBM patients (*P* > 0.05). It was not surprising that Cx43-high recurrent GBM patients exhibited a dismal prognosis (Recurrent GBM), because recurrent GBMs are often refractory to TMZ (5). Similar results were found in multiple GBM datasets (Supplemental Fig. S3A and S4A). To compare Cx43 with other connexins, we performed Cox univariate analyses, which yield a hazard ratio (HR) that determines chance of death (HR > 1 indicates high risk of death). Consistent with the results of Kaplan-Meier analyses (**Fig. 2A**), Cx43-high patients had considerably high HRs in the group of All GBM, MGMT–/TMZ–, and Recurrent GBM. In contrast, most of other connexins failed to displayed a notably high risk of death in all three groups (**Fig. 2B** and Supplemental Fig. S3B and S4B). Cx43 is therefore the only connexin that correlates with poor prognosis of MGMT-deficient GBM.

**Fig. 2.**
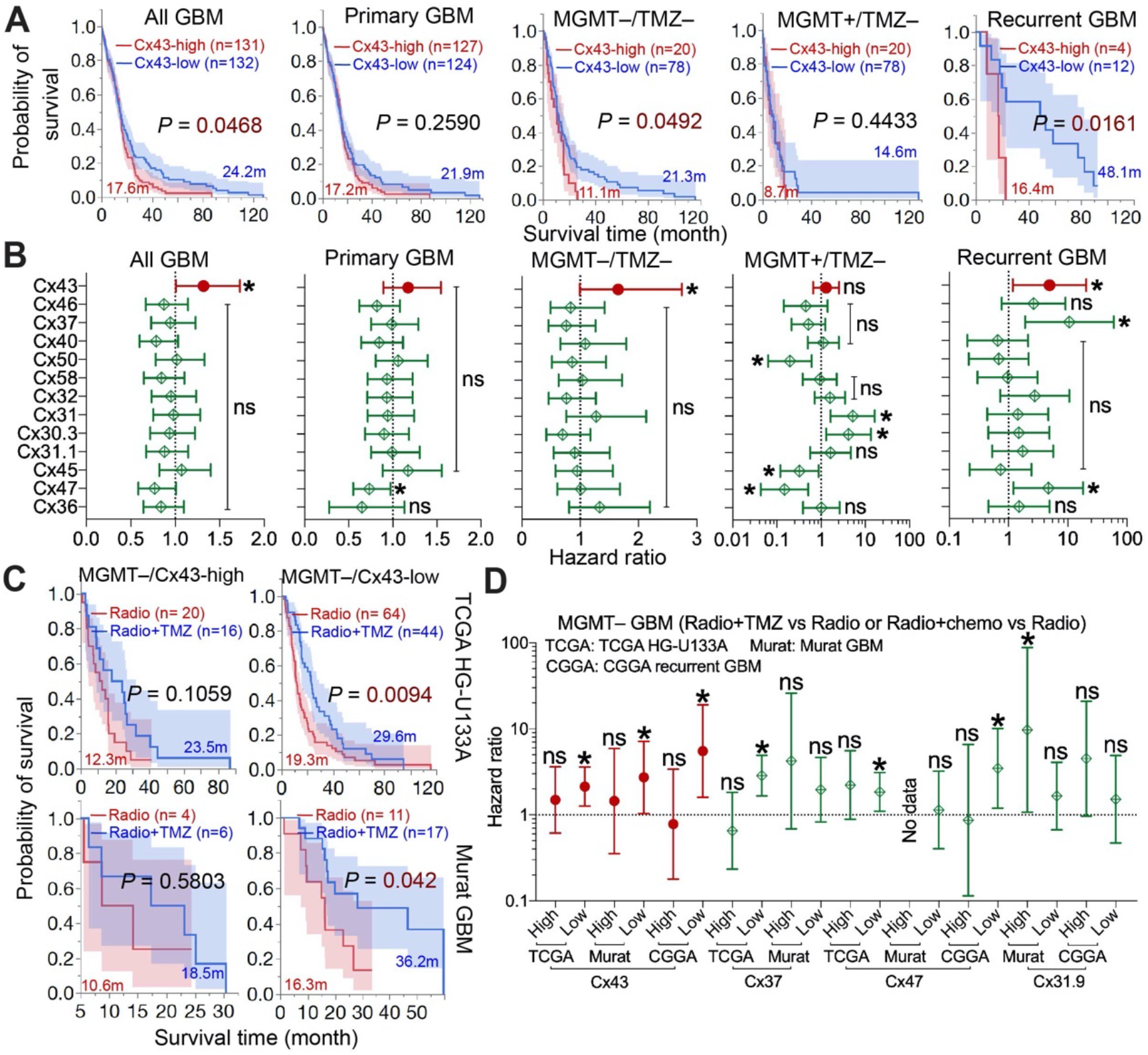
Cx43, but not other connexins, correlates with GBM poor prognosis and chemoresistance. GBM datasets were retrieved from cBioPortal, GlioVis, or CGGA data portal. Immunostaining results of high-grade glioma were obtained from the Human Protein Atlas. (**A**) Kaplan-Meier analysis in the TCGA HG-U133A microarray dataset. Patients were divided into Cx43-high (red; top 25 percentile) or Cx43-low (blue; bottom 25 or 75 percentile) based upon Cx43 mRNA levels in primary, secondary, and recurrent GBM (All GBM), primary GBM only (Primary GBM), MGMT-deficient/TMZ-untreated primary GBM (MGMT–/TMZ–), MGMT-expressing/TMZ-untreated primary GBM (MGMT–/TMZ–), or recurrent GBM only (Recurrent GBM). Case number (n), mean survival time in months (m), and log-rank *P* values are shown. Red or blue shadows represent 95% confidence interval of Cx43-high or Cx43-low group, respectively. (**B**) Cox univariate analysis in the TCGA HG-U133A microarray dataset. The Cox univariate analysis employs the Cox proportional hazards model to yield a hazard ratio that indicates risk levels of death in patients with high levels of connexins compared to those with low levels. The resulting *P* value determines significance of hazard ratio. Cx43 is highlighted in red. (**C**) Kaplan Meier analysis in TCGA HG-U133A and Murat GBM. MGMT-deficient primary GBMs were divided into Cx43-high or Cx43-low group as described above. Patients treated with radiation alone (Radio; red) were compared to patients treated with both radiation and TMZ (Radio+TMZ; blue). (**D**) Cox univariate analysis in TCGA HG-U133A, Murat GBM, and CGGA recurrent GBM. MGMT-deficient primary GBMs or recurrent GBMs were divided into Cx43-high or Cx43-low group. One-Way ANOVA was used to determine statistical significance. *: *P* < 0.05. ****: *P* < 0.0001. ns: not significant.

Previous research has demonstrated that TMZ improves prognosis of GBM patients when used in combination with radiation (6). To determine how connexins contribute to this treatment regime, MGMT-deficient GBM patients treated with radiation (Radio) were compared to patients treated with radiation and TMZ (Radio+TMZ) or radiation and chemo (Radio+chemo) (**Fig. 2C** and Supplemental Figure S5). While the addition of TMZ or chemo did increase the survival of both Cx43-high and Cx43-low patients, there was no statistically significant difference between these treatments in the Cx43-high group in three GBM datasets (*P* > 0.05), suggesting that Cx43-high patients are resistant to TMZ. Of note, levels of Cx37, Cx47, or Cx31.9 did not consistently correspond to the risk of death in three datasets (**Fig. 2D**). Together, our results demonstrate that Cx43 is expressed at the highest level among all connexins in GBM and contributes to chemoresistance as well as poor prognosis of MGMT-deficient GBMs.

### Cx43 confers resistance to TMZ through activating PI3K

Next, we explored how Cx43 confers TMZ resistance. We have previously shown that the Cx43 peptide inhibitor αCT1 inactivates PI3K (21), leading us to hypothesize that Cx43 activates PI3K to induce TMZ resistance. To test this hypothesis, we treated Cx43-high/TMZ-resistant U87MG cells with TMZ or αCT1. αCT1 blocked phosphorylation of Cx43 at serine 368 (**Fig. 3A**, pCx43-S368), a phosphorylation site critical for Cx43 activity (27). As expected, αCT1 induced a 5-fold decrease of the phosphorylated form of AKT serine/threonine kinase (AKT; **Fig. 3A**, pAKT-S473) indicative of a strong inhibition of PI3K. Previous research (28, 29) has suggested that Cx43 regulates the activity of the mitogen-activated protein kinase (MAPK) pathway, including the RAF proto-oncogene serine/threonine-protein kinase (RAF)/extracellular-signal-regulated kinase (ERK) cascade and the SRC proto-oncogene non-receptor tyrosine kinase (SRC) pathway. αCT1 modestly reduced levels of pcRAF-S338, pERK-T202/T204, or pSRC-T416. Hence, αCT1 influences the activity of multiple signaling pathways. The Cx43-induced activation of PI3K was further verified by the knockdown of Cx43 using a short hairpin RNA (shRNA) because the Cx43 shRNA not only drastically decreased levels of Cx43 and pCx43-S368, but also remarkably mitigated PI3K activity in U87MG cells but not in Cx43-low/TMZ-sensitive A172 cells (**Fig. 3B**). Through reanalyzing data from our previous work (21, 30), we detected a strong correlation between Cx43 and pAKT-S473 in six MGMT-deficient GBM cell lines (**Fig. 3C** and Supplemental Table S2). A positive trend was also found between levels of Cx43 mRNA and pAKT-S473 or pAKT-T308 in 37 MGMT-deficient GBM patients in the TCGA dataset (**Fig. 3D**). Other connexins, however, failed to show statistically significant correlations with either pAKT-S473 (**Fig. 3E**) or pAKT-T308 (**Fig. 3F**).

**Fig. 3.**
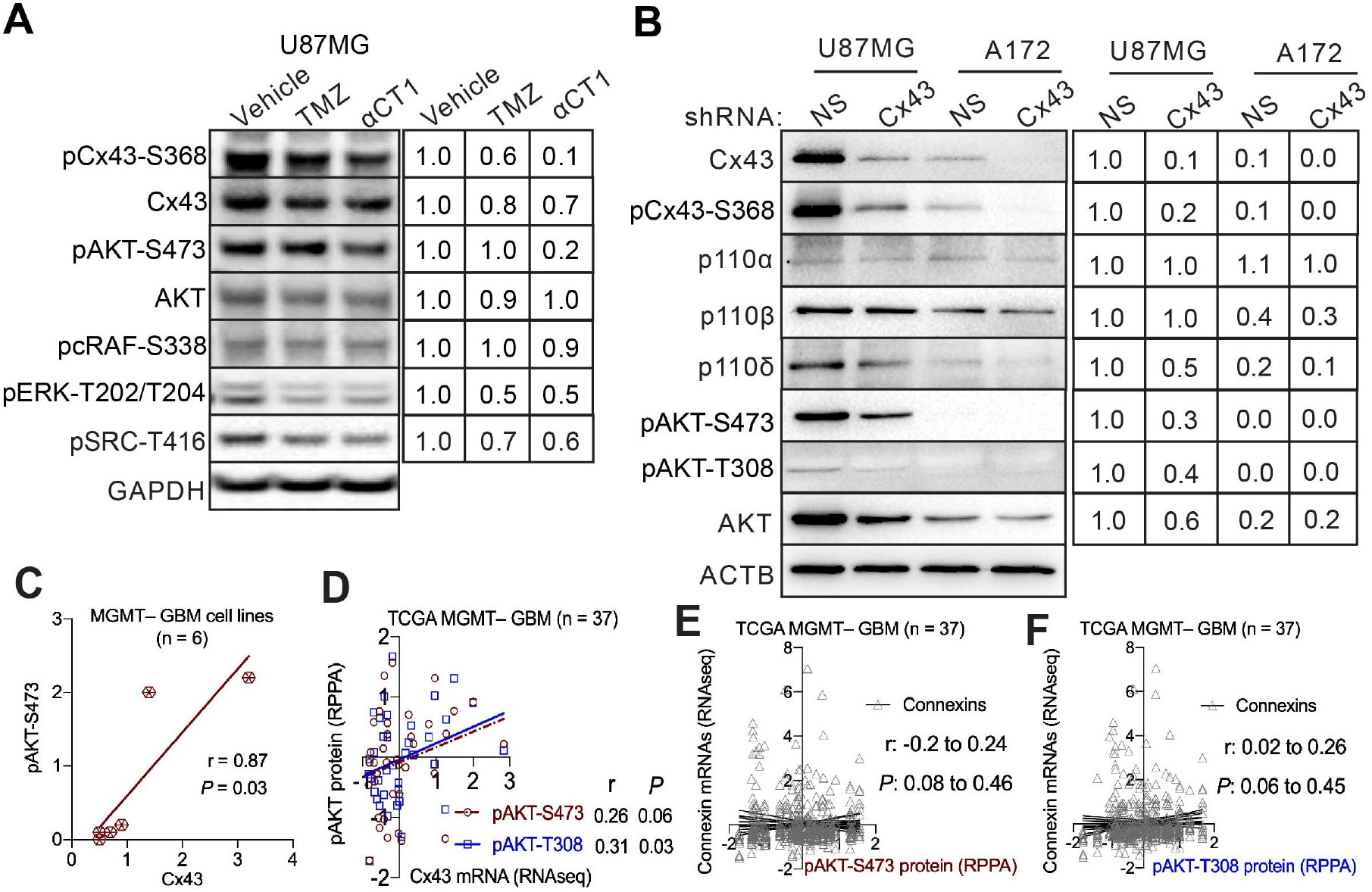
Cx43 blockade inactivates PI3K. (**A**) Signaling pathways affected by αCT1. U87MG cells were treated with 100 μM αCT1 or 50 μM TMZ for 4 days. pAKT-S473, pcRAF-S338, pERK-T202/T204), and pSRC-T416 were analyzed using immunoblotting. Glyceraldehyde 3-phosphate dehydrogenase (GAPDH) was the loading control. Band intensities were quantified using Image J. Vehicle was set as 1.0 and each treatment was normalized to the vehicle. (**B**) PI3K signaling upon depletion of Cx43. U87MG and A172 cells were treated with a non-silencing short hairpin RNA (NS shRNA) or a Cx43 shRNA. U87MG cells treated with NS shRNA were set as 1.0. β-actin (ACTB) was the loading control. Pearson coefficient correlation analysis between protein levels of Cx43 and pAKT-S473 in 6 MGMT-deficient GBM cell lines (**C**), mRNA levels of Cx43 and protein levels of pAKT-S473 or pAKT-T308 in MGMT-deficient patients (**D**), or mRNA levels of connexins and protein levels of pAKT-S473 (**E**) and pAKT-T308 (**F**) in MGMT-deficient GBMs. The Pearson correlation coefficient (r) and *P* value that determines statistical significance of the coefficient are shown. Cell line data were retrieved from our previous studies (21, 30). RNA sequencing (RNAseq) data and results of reverse phase protein array (RPPA) were retrieved from the TCGA database.

To determine whether PI3K is required for Cx43-induced TMZ resistance, we overexpressed PIK3CA-E545K, a PI3K mutant that constitutively activates PI3K, in U87MG cells (**Fig. 4A**). PIK3CA-E545K counterbalanced the growth inhibition induced by TMZ or by a combination of TMZ and αCT1 (**Fig. 4B**). This counteraction was not seen in U87MG cells expressing an active mutant of ERK (ERK2-L73PS151D; **Fig. 4C**) or SRC (SRC-Y527F; **Fig. 4D**). These results suggest that, while Cx43 activates multiple signaling pathways such as PI3K, ERK, or SRC, only the activation of PI3K is important for Cx43 to induce TMZ resistance.

**Fig. 4.**
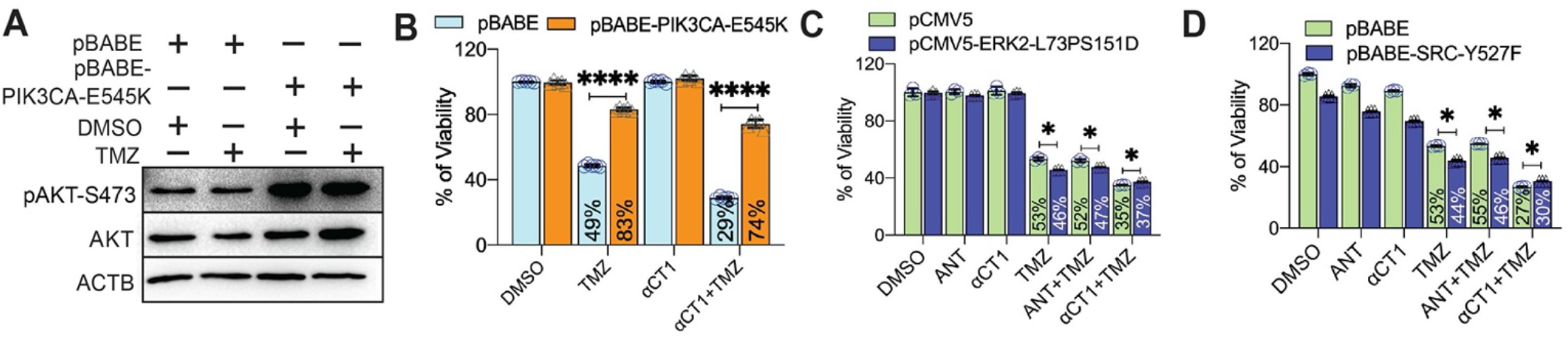
Activation of PI3K, but not ERK or SRC, reverse growth inhibition induced by αCT1/TMZ. (**A**) Expression of PIK3CA-E545K (an active PI3K mutant). U87MG cells were transfected with pBABE or pBABE-PIK3CA-E545K encoding PIK3CA-E545K followed by the treatment of 100 μM TMZ. Dimethyl Sulfoxide (DMSO) is the vehicle control. (**B**) The effect of PIK3CA-E545K on the αCT1/TMZ-induced growth inhibition. Transfected cells were treated with a combination of 100 μM αCT1 and/or 100 μM TMZ for 6 days. Cell viability was measured using the MTS viability assay. Percentages of viability were obtained by normalizing the MTS readings of treatment groups to that of DMSO. (**C**) The effect of ERK2-L73PS151D on the αCT1/TMZ-induced growth inhibition. U87MG cells were transfected with pCMV5 or pCMV5-ERK2-L73PS151D (encoding an active ERK2 mutant) followed by the treatment of αCT1 or antennapedia peptide (ANT; the control peptide for αCT1) and/or TMZ. (**D**) The effect of SRC-Y527F on the αCT1/TMZ-induced growth inhibition. U87MG cells were transfected with pBABE or pBABE-SRC-Y527F (encoding an active SRC mutant) followed by the treatment of αCT1 or ANT and/or TMZ. Student *t* test was used to determine statistical significance. *: *P* < 0.05; ****: *P* < 0.0001.

### Cx43 activates PI3K through selectively binding to the PI3K catalytic subunit β

Because the Cx43-CT regulates the activity of Cx43-channels (31), it is possible that small molecules such as ATP or glutamate released from Cx43-channels activate PI3K in GBM cells as they do in astrocytes (32). To test this possibility, we treated U87MG cells with Gap27, a Cx43 peptide inhibitor that targets the second extracellular loop of Cx43 and blocks Cx43-channels (33). Gap27, however, did not attenuate PI3K activity (**Fig. 5A**). Moreover, levels of ATP or glutamate in culture media either elevated or remained unchanged in αCT1-treated cells (**Fig. 5B-D**), consistent with the dephosphorylation of Cx43 at S368 by αCT1 (**Fig. 3A**), which enhances the permeability of Cx43 hemichannels (34). ATP or glutamate levels remained unchanged in cells (**Fig. 5E-F**). Our results suggest that Cx43-channels are dispensable for PI3K activation in GBM cells.

**Fig. 5.**
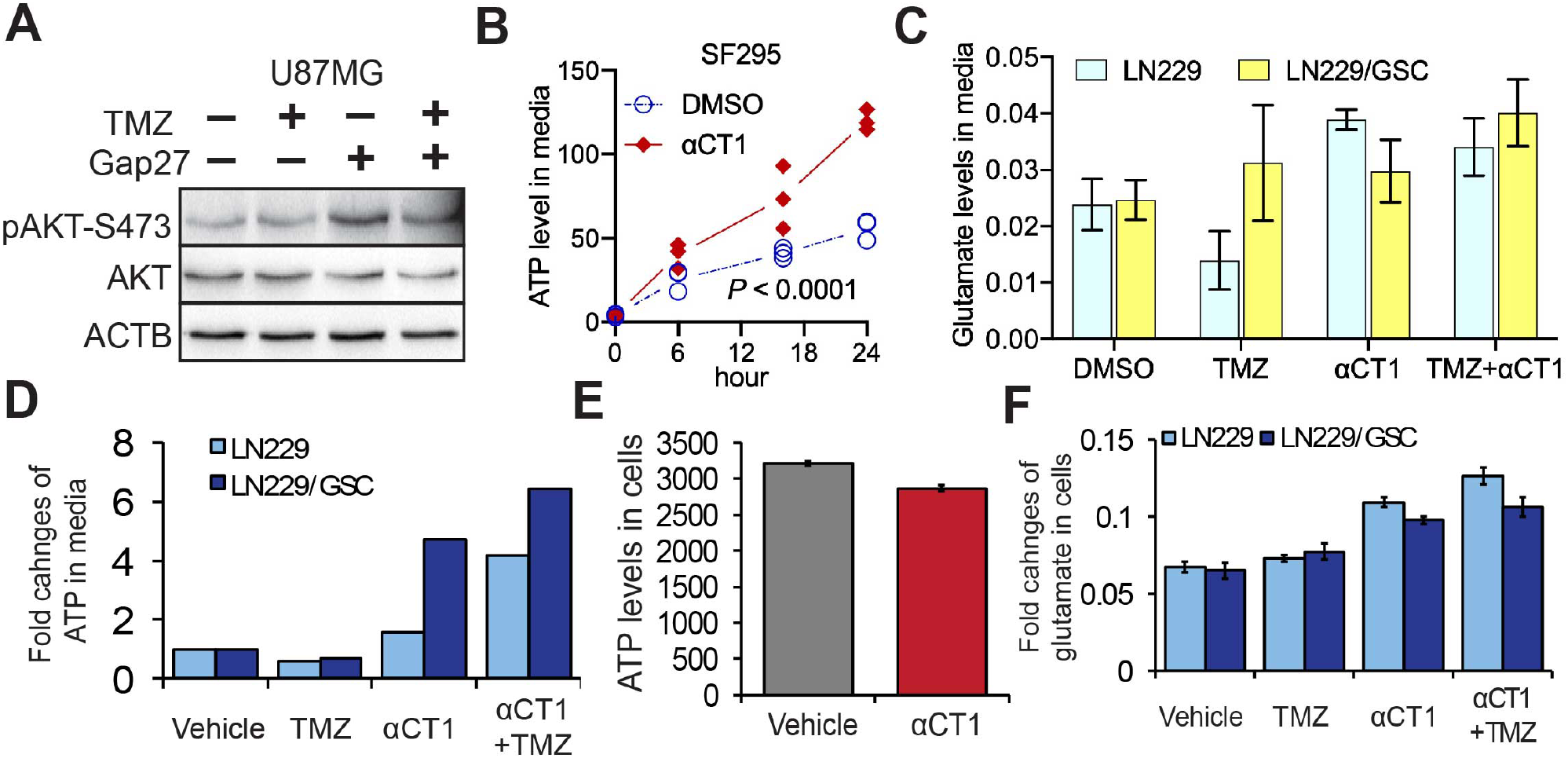
αCT1 does not change Cx43 channel activity in GBM. (**A**) The effect of Gap27 on PI3K signaling. U87MG cells were treated with 100 μM TMZ or 100 μM Gap27. (**B**) ATP release from Cx43-high/MGMT-deficient/TMZ-resistant SF295 cells. Cells were treated with 100 μM αCT1. Culture media were collected at different time points. ATP was measured using a colorimetric assay. Oneway ANOVA was used to determine statistical significance. (**C**) Glutamate release in Cx43-low/MGMT-deficient/TMZ-sensitive LN229 or Cx43-high/MGMT-deficient/TMZ-resistant LN229/GSC cells. Cells were treated with 100 μM TMZ and/or 100 μM αCT1. Glutamate in culture media was determined using a colorimetric assay. (**D**) ATP release in LN229 and LN229/GSC cells. (**E**) ATP within SF295 cells. (**F**) Glutamate within LN229 and LN229/GSC cells. GSC: glioblastoma stem cells.

Cx43-CT interacts with certain signaling molecules (28). It is likely that Cx43 binds to PI3K catalytic subunits to activate PI3K. The Class I PI3K family consists of four highly homologous catalytic subunits: PI3K catalytic subunits α, β, δ, and γ (PIK3CA, PIK3CB, PIK3CD, and PIK3CG) encoding p110α, p110β, p110δ, and p110γ, respectively (35). Our previous work has demonstrated that PI3K catalytic subunits play different roles in GBM cell survival, with p110β being the most dominant isoform in GBM (30). To determine whether PI3K catalytic subunits also function divergently in Cx43-induced PI3K activation, we reanalyzed protein expression data in six MGMT-deficient GBM cell lines (Supplemental Table S2). Levels of Cx43 protein showed a positive and statistically significant correlation with those of p110β, but not other p110s or the regulatory subunit p85 (**Fig. 6A**). mRNA levels of Cx43 also positively corresponded with those of PIK3CB, but not other PI3K subunits, in 89 MGMT-deficient GBM patients in the TGCA RNAseq dataset (**Fig. 6B**). In the same dataset, PIK3CB displayed no or negative correlation with the 21 other connexin family members, except Cx31 (**Fig. 6C**). Such a positive correlation between Cx43 and PIK3CB was recapitulated in multiple GBM datasets (Supplement Fig. S6) and further verified by the finding that high levels of pAKT-S473 or p110β, but not other p110s, correlated with low TMZ sensitivity indicated by the increase of TMZ IC50s (**Fig. 6D-E**). To further probe the molecular details of Cx43-induced PI3K activation, we monitored protein-protein interactions between Cx43 and p110 proteins. Cx43 was co-precipitated with p110β (**Fig. 6F**), but not with p110α or p110δ (**Fig. 6G-H**), demonstrating a selective binding of Cx43 to p110β. We did not examine p110γ because p110γ is not detectable in GBM (30). To determine whether αCT1 binds to Cx43 and/or p110β, we treated U87MG cell lysates with αCT1 and found that αCT1 was pulled down together with p110β and Cx43 (**Fig. 6I**). In the presence of αCT1, more p110β was found in the Cx43 precipitate. This might be because the Cx43 antibody is able to precipitate αCT1/Cx43/p110β (or αCT1/p110β) and Cx43/p110β protein complexes. Taken together, we have, for the first time, defined a novel non-channel activity of Cx43, through which p110β is selectively bound and activated in GBM.

**Fig. 6.**
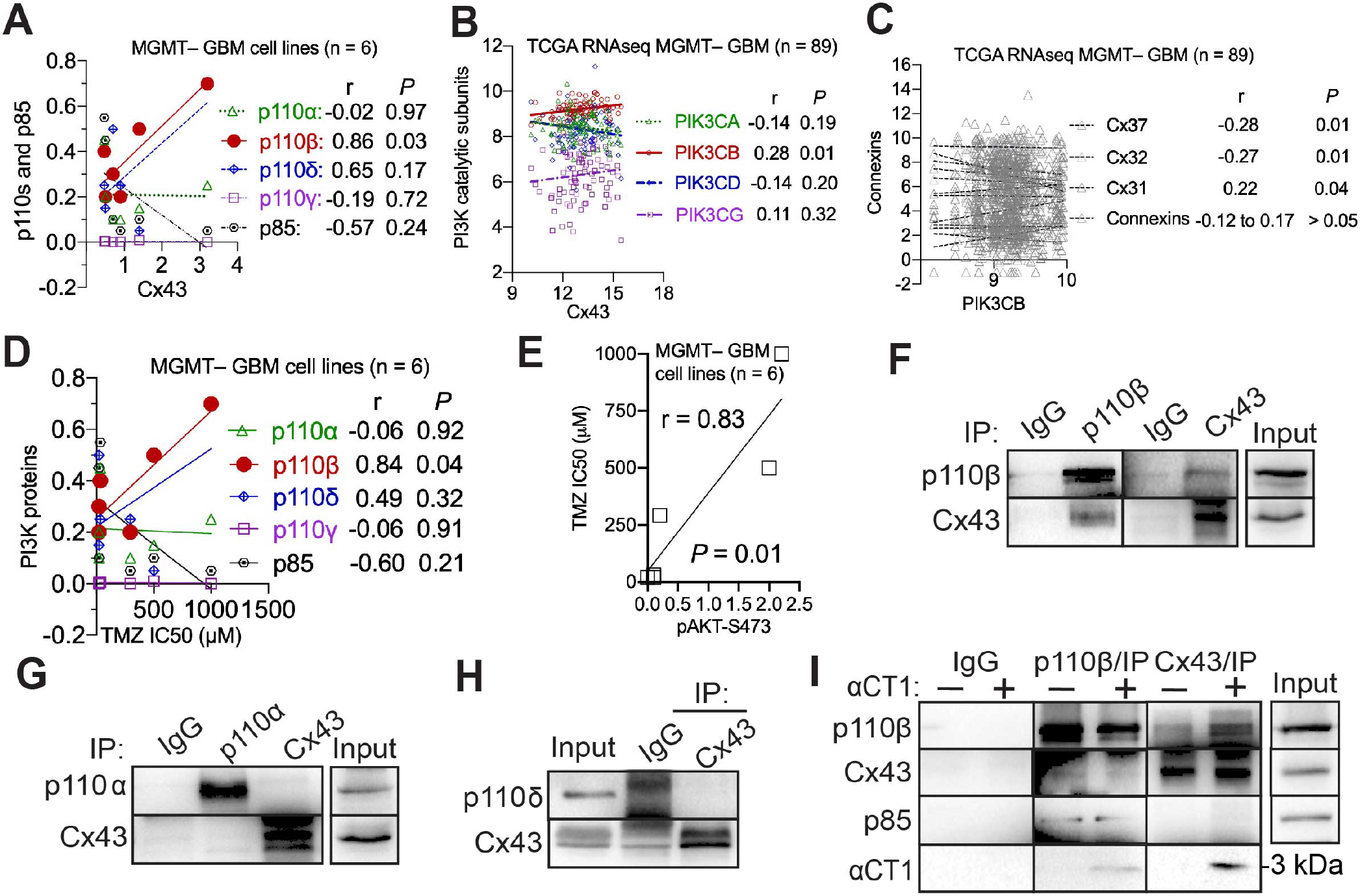
Cx43 activates PI3K through selectively binding to the PI3K catalytic subunit β. Pearson coefficient Correlation between protein levels of Cx43 and PI3K catalytic subunits in 6 MGMTdeficient GBM cell lines (**A**), mRNA levels of Cx43 and PI3K catalytic subunits in MGMT-deficient GBM patients (**B**), mRNA levels of PIK3CB and connexins in MGMT-deficient GBM patients (**C**), protein levels of p110 proteins and IC50s of TMZ in 6 MGMT-deficient GBM cell lines (**D**), or protein levels of pAKT-S473 and IC50s of TMZ in 6 MGMT-deficient GBM cell lines (**E**). Cell line data were retrieved from our previous studies (21, 30). RNAseq data were retrieved from the TCGA database. The Pearson correlation coefficient r and corresponding *p* are shown. Co-immunoprecipitation of Cx43 and p110β (**F**), p110α (**G**), or p110δ (**H**) in U87MG cells. (**I**) Co-immunoprecipitation of Cx43 and p110β in U87MG cell lysates treated with 100 μM αCT1. αCT1 is about 3 kDa and recognized by the Cx43 antibody. IP: immunoprecipitation. Rabbit IgG was used as the control.

### A combination of αCT1 and p110β-selective inhibitors overcomes TMZ resistance

αCT1 alone increases the sensitivity of LN229/GSC xenograft tumors to TMZ (21); however, the short half-life of αCT1 demands high concentrations and repeated drug delivery, which may limit its therapeutic potential. Prompted by the above results, we tested the combination of αCT1 and p110β-selective inhibitors in cultured cells and in mice. To achieve a synergistic therapeutic effect of multiple drugs, we optimized the dose of each individual drug in U87MG cells. By varying doses of TMZ or a p110β-selective inhibitor TGX-221, we found that the double combination of 50 μM TMZ and 20 μM TGX-221 did not significantly inhibit the viability of U87MG cells (Supplemental Fig. S7A-B). However, the addition of αCT1 greatly increased the cytotoxic effect of the TMZ/TGX-221 double combination (Supplemental Fig. S7C). We next employed a coefficient of drug interaction (CDI) analysis, a method that has been used to measure drug synergy (36–39). CDI < 1 indicates a synergistic effect; CDI = 1 means an additive drug effect; CDI > 1 refers to an antagonistic effect. 2.5 to 10 μM αCT1 only yielded an additive effect together with TMZ/TGX-221, whereas 12.5 to 50 μM αCT1 synergistically blocked cell growth (Supplemental Fig. S7D).

Based on these results, 30 μM αCT1, 20 μM TGX-221, and 50 μM TMZ was used in a triple combination named αCT1/TGX/TMZ combo. The αCT1/TGX/TMZ combo synergistically reduced the viability of MGMT-deficient/TMZ-resistant SF295, VTC-103, and VTC-003 cells that express high levels of Cx43 and p110β (**Fig. 7A** and Supplemental Fig. S8A) (21, 30). Notably, VTC-103, VTC-003, and other VTC lines described hereafter were derived from freshly dissected GBM tumors (21, 30). CDIs of the αCT1/TGX/TMZ combo were significantly lower than those of double combinations (**Fig. 7B** and Supplemental Fig. S8B). This synergistic effect was, however, not found in MGMT-deficient/TMZ-sensitive LN229 and A172 or MGMT-deficient/TMZ-resistant VTC-001, VTC-005, and VTC-004 (**Fig. 7C-D** and Supplemental Fig. S8 and S9) whose levels of Cx43 and p110β are low (21, 30). The αCT1/TGX/TMZ combo activated apoptosis in VTC-103 cells (**Fig. 7E**), coinciding with the drastic decrease of cell growth (**Fig. 7A**), whereas apoptosis was not induced in VTC-001 cells (**Fig. 7E**). To verify our *in vitro* studies *in vivo*, we treated mice bearing SF295 xenograft tumors with 7.5 mg/kg TMZ and 20 mg/kg TGX-221 through intraperitoneal injection in conjunction with 32.6 μg of αCT1 per tumor through intratumoral injection. The αCT1/TGX/TMZ combo (red line) stopped tumor growth (**Fig. 7F**, *P* < 0.05), whereas double combinations exhibited limited to no inhibition. Based on tumor volumes on the last day of treatment, the CDI of the triple combo was approximately 0.22, confirming a strong synergy amongst αCT1, TGX-221, and TMZ *in vivo*. To verify that the synergistic cytotoxicity is due to the blockade of Cx43/p110β, we knocked down Cx43 and individual PI3K catalytic subunits using shRNAs. Depletion of p110β, but not p110α or p110δ, blocked the growth of SF295 cells (**Fig. 7G**) and only the combination of PIK3CB shRNA, Cx43 shRNA, and TMZ yielded synergistic inhibition of cell viability (**Fig. 7H**).

**Fig. 7.**
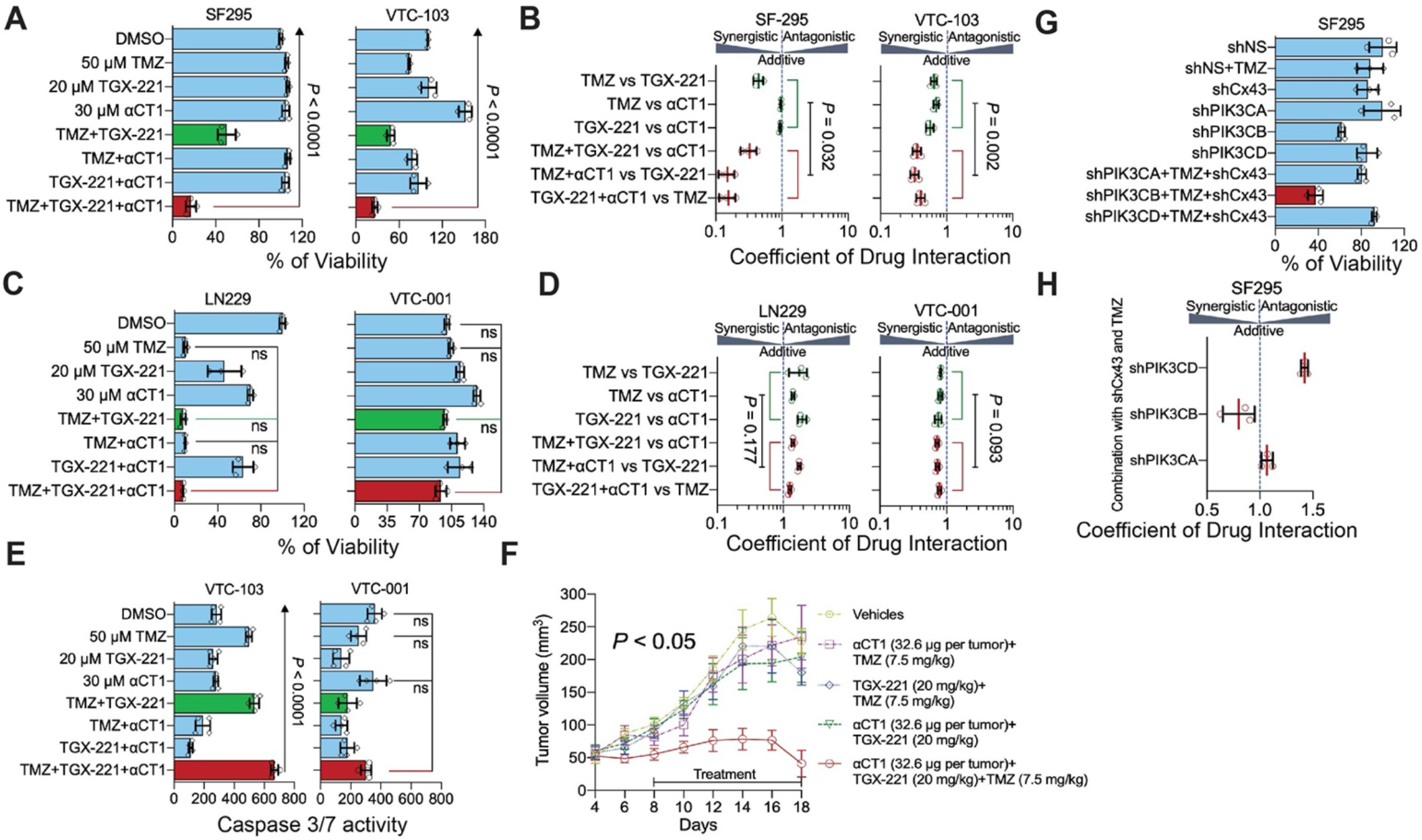
A combination of αCT1 and TGX-221 overcomes TMZ resistance *in vitro* and *in vivo*. (**A**) The effect of the αCT1/TGX-221/TMZ combo in Cx43/p110β-high/MGMT-deficient/TMZ-resistant SF295 and VTC-103 cells. Cells were treated with 50 μM TMZ, 20 μM TGX-221, and/or 30 μM αCT1 including single agents, double combinations and the αCT1/TGX-221/TMZ combo. This scheme has been repeated in experiments presented hereafter. Cell viability was determined using the MTS viability assay. Percentages of cell viability were obtained by normalizing the MTS readings of treatment groups to that of DMSO group. (**B**) Coefficient of drug interaction (CDI) analysis in SF295 and VTC-103 cells. The drug combination is synergistic if CDI is less than 1, additive if CDI equals to 1, or antagonistic if CDI is more than 1. CDIs of triple combinations (red) were compared to double combinations (green) and statistical significance was determined using the student *t* test. (**C**) The effect of the αCT1/TGX-221/TMZ in Cx43/p110β-low/MGMT-deficient/TMZ-sensitive LN229 and TMZ-resistant VTC-001 cells. (**D**) CDI analyses in LN229 and VTC-001 cells. (**E**) Caspase 3/7 activity in VTC-103 and VTC-001 cells. The activity of cleaved caspase 3/7 (active) was determined using a luminescence assay. Shown are luminescence readings. (**F**) The effect of αCT1/TGX-221/TMZ combo on SF295 xenograft tumors. SF295 cells were subcutaneously injected into immuno-deficient mice. 8 days later, mice were treated with TMZ, TGX-221, or αCT1 through intraperitoneal or intratumoral injection every other day untill day 18. Tumor volumes are shown. (**G**) The effect of shRNA of Cx43 or PI3K catalytic subunits on the TMZ sensitivity of SF295 cells. Cells were transfected with NS shRNA or shRNA of Cx43, PIK3CA, PIK3CB, or PIK3CD followed by the treatment of 50 μM TMZ. Cell viability was determined using the MTS viability assay. Percentages of cell viability were obtained by normalizing the MTS readings of treatment groups to that of shNS group. (**H**) The coefficient of drug interaction (CDI) analyses for **A**. One-way ANOVA or student *t* test was used to determine statistical significance. *: *P* < 0.05; ns: not significant.

To corroborate results from TGX-221, we tested another p110β-selective inhibitor GSK2636771 (abbreviated hereafter as GSK), which has been used in a clinical study (40). αCT1/GSK/TMZ combo entailing 25 μM GSK, 30 μM αCT1, and 50 μM TMZ synergistically blocked the viability of VTC-103 cells (**Fig. 8A-B**) and U87MG cells (Supplemental Fig. S10), but not the viability of LN229 cells (**Fig. 8C-D**). αCT1/GSK/TMZ has achieved the same synergistic inhibition of GBM cell viability as the αCT1/TGX/TMZ combo. To determine the toxicity of these combinations on normal cells, we treated astrocytes with αCT1/TGX/TMZ or αCT1/GSK/TMZ. These drug combinations did not increase TMZ alone-induced growth inhibition in astrocytes (**Fig. 8E**), suggesting that addition of αCT1 and p110β-selective inhibitors does not exacerbate non-selective toxicity of TMZ to the normal brain. Collectively, our results demonstrate that simultaneously targeting Cx43 and p110β diminishes TMZ resistance.

**Fig. 8.**
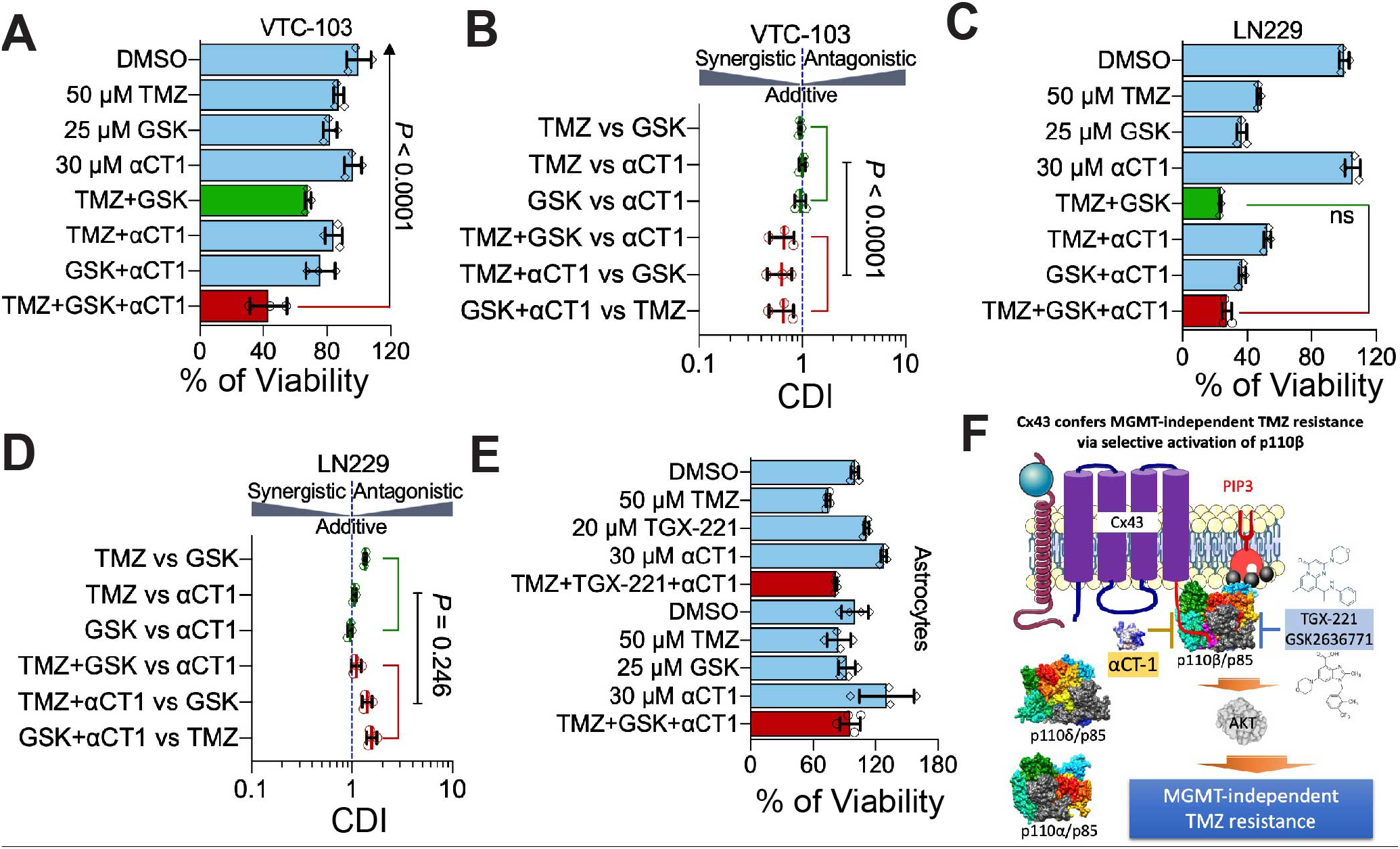
A combination of αCT1 and GSK2636771 overcomes TMZ resistance. (**A**) The effect of the αCT1/GSK/TMZ combo in VTC-103 cells. Cells were treated with 50 μM TMZ, 25 μM GSK2636771, and/or 30 μM αCT1 including single agents, double combinations and the αCT1/GSK/TMZ combo. (**B**) CDI analyses in VTC-103 cells. (**C**) The effect of the αCT1/GSK/TMZ combo in LN229 cells. (**D**) CDI analyses in LN229 cells. (**E**) The effect of the αCT1/TGX-221/TMZ or αCT1/GSK/TMZ combo in astrocytes. (F) A model illustrating the mechanism of Cx43-induced MGMT-independent TMZ resistance and the model of action of the triple combination. One-way ANOVA or student *t* test was used to determine statistical significance. *: *P* < 0.05; ns: not significant.

## Discussion

In this report, we have identified the molecular details underlying Cx43-induced MGMT-independent TMZ resistance. As illustrated in a model proposed in **Fig. 8F**, Cx43-CT binds to p110β/p85 signaling complex upon receiving signals from extracellular cues (i.e. growth factors). This selective binding brings the p110β/p85 signaling complex to the membrane and subsequently activates AKT. Activated PI3K/AKT signaling renders GBM cells resistant to TMZ, which is independent of MGMT. This model not only explains how a gap junction protein regulates chemoresistance through its non-channel activity, but also provides a strong rationale for developing combinational therapies to overcome TMZ resistance. Indeed, our results shown in **Figs. 7–8** indicates that αCT1, a Cx43-CT mimetic peptide that likely blocks interactions between Cx43-CT and p110β, works synergistically together with p110β kinase inhibitors (directly blocking kinase activity) in overcoming TMZ resistance.

Prior studies report that approximately 20-60% of GBM patients express Cx43 mRNA and protein at high levels (15). In light of the fact that 45% of GBM patients express no MGMT (6, 7), there should be 10% (20% x 50%) to 30% (60% x 50%) of patients that are MGMT-deficient and express high levels of Cx43. Congruent with this expectation, we have found that 16.7% of MGMT-deficient GBM patients express high levels of Cx43 (21). That being said, around 20% of Cx43-high GBM patients may be refractory to TMZ treatment in the clinic. Therefore, the combinational treatment developed herein will benefit MGMT-deficient/TMZ-resistant patients expressing high levels of Cx43, thereby having an important impact on future therapeutic intervention. Previous work has also revealed that, with the exception of Cx43, overexpression or inhibition of Cx30, Cx32, Cx26, or Cx46 also blocks the growth of rat or human glioma cells (41–48). However, contradictory to these results, other studies show that Cx30 and Cx32 have no effect on glioma growth (44, 49, 50). In line with the fact that Cx43 levels are much higher than other connexins in GBM and the finding that Cx43 controls chemoresistance, this connexin is therefore the prime therapeutic target for GBM.

Cx43 has long been considered as a tumor suppressor for glioma because overexpression of Cx43 leads to remarkable growth inhibition (51) and levels of Cx43 mRNA and protein inversely correlate with the aggressiveness of glioma (52). However, drawbacks in these studies have made the tumor suppressive activity of Cx43 questionable. For example, while ectopically expressing Cx43 does inhibit tumor cell growth, it is unclear whether the loss of endogenous Cx43 in normal glia cells promotes gliomagenesis as other tumor suppressors do, namely p53 and NF-1. Nonetheless, it is possible that gap junction intercellular communication controlled by Cx43 is GBM suppressive because loss of this communication promotes oncogene-induced transformation (53). In contrast to these studies, we have established a tumor-promoting role of Cx43 in GBM. Cx43, whose mRNA levels are the highest among all connexins, not only correlates with GBM prognosis and chemoresistance, but also activates PI3K independent of Cx43-channels to induce TMZ resistance. Therefore, it is likely that Cx43 has multifaceted roles in GBM: Cx43-channels inhibit GBM formation, whereas the Cx43 CT confers chemoresistance through activating PI3K, which is independent of Cx43 channel function, during GBM progression.

## Methods

### Reagents

TMZ (AbMole BioScience), GSK2636771 (AdooQ Bioscience), TGX-221 (AdooQ Bioscience) were reconstituted in dimethyl sulfoxide (DMSO) at a concentration of 50-80 mM. αCT1 and Gap27 were purchased from LifeTein, LLC. Lyophilized peptide was reconstituted in 1x PBS (137 mM NaCl, 2.7 mM KCl, 10 mM Na_2_HPO_4_, and 1.8 mM KH_2_PO_4_) at a concentration of 5 or 10 mM. Puromycin was purchased from Millipore-Sigma and dissolved in sterile water at a concentration of 5 mg/ml. All chemicals were aliquoted (to avoid repeated freeze/thaw cycles that decrease drug activity) and stored at −80 °C.

### Cell culture

GBM cell lines, primary GBM cells, glioblastoma stem cells (GSCs), and human astrocytes were cultured as previously described (54). In brief, GBM cell lines A172, SF-295, LN229, and U87MG were maintained in Dulbecco’s Modified Eagle Medium (DMEM, Life Technologies) supplemented with 10% EquaFETAL^®^ bovine serum (Atlas Biologicals, Inc.) and 100 μg/ml streptomycin and 100 IU/ml penicillin (Gibco). Primary cells VTC-001, VTC-003, VTC-04, VTC-005, and VTC-103 were cultured in DMEM supplemented with 15% fetal bovine serum (Peak Serum, Inc.) and penicillin/streptomycin. Normal human astrocytes were cultured in MCDB-131 Medium (Sigma) containing 3% fetal bovine serum (Peak Serum, Inc.), 10 X G-5 Supplement (Gibco), and penicillin/streptomycin. Cell lines have been authenticated by the ATCC authentication service utilizing Short Tandem Repeat (STR) profiling. Primary GBM cells were kept at low passages (no more than 10).

### Analysis of online databases

GBM gene expression datasets (cDNA microarrays or RNA sequencing) or the data of reverse phase protein assay (RPPA) associated with corresponding clinical information and protein immunostaining data of human tissues are downloaded from the following websites: (1) The Cancer Gene Atlas (TCGA) datasets: https://www.cbioportal.org and https://gliovis.shinyapps.io/GlioVis/; (2) Gravendeel, Rembrandt, Lee Y, and Murat GBM: https://gliovis.shinyapps.io/GlioVis/; (3) The China Glioma Gene Atlas (CGGA) datasets: https://gliovis.shinyapps.io/GlioVis/; (4) GBM cell lines from the Cancer Dependency Map (DepMap): https://depmap.org/portal/; (5) The Human Protein Atlas (THPA): https://www.proteinatlas.org. To analyze gene expression data, arbitrary readings from cDNA microarray or RNAseq were used. mRNA levels of GJ genes were averaged and plotted using the Prism 8 software. For protein immunostaining data, histological images were downloaded and presented. Staining scales of each sample were recorded. Percentage of cases with high levels or not high levels of GJ proteins were plotted using the Prism 8 software.

For survival analysis, patient clinical information was matched to each individual gene such as GJA1. Different populations of GBM patients were sorted based on their clinical information. The Kaplan Meier survival analysis or the Cox hazard proportional model were used to determine the relationship between gene expression levels and patient survival. The JMP Pro 15 software was used to perform these analyses. For Kaplan Meier survival analysis, log-rank *P* values are shown to indicate statistical significance of patient survival between high and low levels of a given gene. For Cox hazard proportional model, hazard ratios that indicate the risk of death in patients with high levels of a given gene are shown to present a comparison among different GJ genes.

To determine the expression correlation between different genes or proteins, Pearson correlation coefficient was calculated using the Prism 8 software. The Pearson correlation coefficient determines whether expression levels of two genes are correlated with each other (positively or negatively) and whether this correlation is statistically significant or not.

### MTS cell viability assay

Cell viability was determined by the MTS cell viability assay (Promega) as described previously (21, 30, 55–57). In brief, 250 to 1,000 cells were plated in the wells of a 96-well plate based upon the cell growth rate. Because the drug treatment usually takes 6-7 days, fast-growing cells could be over-grown if plated at a high cell density. For αCT1 treatment experiments, we intended to plate cells at a low density to minimize the formation of gap junctions and thus more Cx43-hemichannels will be present. Because the half-life of αCT1 is about 48 hours, cells were replenished with fresh αCT1 every other day, without replenishing other drugs. Cells were treated with vehicle (DMSO) and chemical inhibitors at the indicated doses. After 6 days MTS reagent was added to the cells to a final dilution of 10% and incubated at 37°C for a 4-hour period. At each hour timepoint the absorbance at 490 nm was measured using a FilterMax F3 microplate reader (Molecular Devices, LLC) according to the manufacturer’s instructions. Percent cell viability was obtained by dividing the absorbance of treatment groups to those of untreated and respective vehicle control groups.

### Caspase 3/7 activity assay

Apoptosis was measured using the Caspase-Glo^®^ 3/7 Assay (Promega) based on the manufacturer’s instructions and our previous work (21, 30, 55, 56). In brief, VTC-001 and VTC-103 cells were plated at 1,000 cells/well in 96-well plates and treated with drugs as described for 6 days. After 6 days, 100 μL of Caspase-Glo^®^ reagent was added to each well and incubated at room temp (RT). The luminescence was measured using a FilterMax F3 microplate reader (Molecular Devices, LLC) according to the manufacturer’s instructions. The fold changes of caspase-3/7 activity was defined as the ratio of caspase-3/7 luminescence in the treated cells to that in control cells.

### Immunoblotting

Immunoblotting was performed as described previously (58, 59). In brief, cells were lysed and total protein was quantified using the Bradford Assay (Bio-Rad Laboratories, Inc.) An equal amount of total protein (25-50 μg) of cell lysate was loaded onto a 15% SDS-PAGE gel and separated proteins were transferred onto a PVDF membrane. The resulting protein blot was incubated with antibodies purchased from Cell Signaling Technology (CST), Millipore-Sigma (MS), and SantaCruz Biotechnology (SC). Antibodies were diluted as follows: anti-phospho-Cx43-S368(CST-3511,1:1,000), anti-Cx43(CST-3512, 1:1,000), anti-phospho-AKT-S473(CST-4051,1:1,000), anti-phospho-AKT-T308(CST-4056,1:1,000), anti-AKT(CST-4685,1:1,000), anti-phospho-cRAF-S338(CST-9247,1:1,000), anti-phospho-ERK-T202/T204(CST-4377,1:1,000), anti-phospho-SRC-T4160(CST-2101,1:1,000), anti-p110α(CST-4249,1:1,000), anti-p110β(CST-3011,1:1,000), anti-p110δ(CST-34050,1:1,000), anti-p85(CST-4292,1:1,000), anti-β-actin(MS-A3854,1:50,000), and anti-GAPDH(SC-25778,1:1,000). Protein bands were visualized using a ChemiDoc MP System (Bio-Rad Laboratories, Inc.) and further quantified using Image J software. The relative level of protein is defined as the ratio of band intensity of target protein to that of β-actin or GAPDH.

### Co-immunoprecipitation

Co-immunoprecipitation was performed as previously described (59). U87MG cells were cultured under normal cell culturing conditions and collected at 80% confluency. Cell pellets of approximately 250 μL volume were flash frozen. Pellets were then lysed in lysis buffer containing 20 mM HEPES pH 6.8, 140 mM NaCl, 2.5 mM MgCl2, 2.5 mM CaCl2, 1% NP40, 0.5% sodium deoxycholate, protease inhibitor (Millipore-Sigma, MS), and phosphatase inhibitors (MS). Total protein lysates were divided equally for each IP with input and IgG controls. Samples were incubated with primary antibodies O/N at 4°C on a rotator. Antibodies were diluted as follows: anti-Cx43(MS-C6219,1:50), anti-p110α(CST-4249,1:25), anti-p110β(CST-3011,1:25), anti-p110δ(CST-34050,1:25). All antibodies were from Rabbit, thus Rabbit IgG (SC-2027,1:400) was used as a control. Samples were then incubated at RT for 1 hour with Protein G Dynabeads™ (Thermo-Fisher) on a rotator. Beads were then precipitated using a magnet and the supernatant removed. Protein-bead complexes were washed 3X with lysis buffer and incubated at RT on a rotator for 10 minutes and pulled down on the magnet each time. After the final wash and supernatant removal, the beads were mixed with 2X SDS loading buffer and 1M DTT and boiled at 95°C for 10 minutes and then vortexed. After boiling, samples were placed on the magnet to remove the beads and the precipitated proteins were ran on a 15% SDS-PAGE gel at equal volume across samples including the IgG and input controls.

### Gene knockdown

Knockdown of Cx43 or PI3K genes was described previously (30). Short hairpin (sh) RNA of Cx43 (TRCN0000059773), previously verified was purchased from Millipore-Sigma. shRNAs previously verified for PI3K genes were purchased from Thermo-Fisher Scientific (PIK3CA: RHS4844-101656239; PIK3CB:RHS4884-10165656350; PIK3CD:RHS4884-101655755). 1×10^6^ HEK-293T cells were transfected with 2 μg of plasmid DNA containing each shRNA along with packaging plasmids pMD2.g and psPax2 using Effectene^®^ Transfection Reagent (QIAGEN) to yield 5 mL of supernatant containing lentivirus. 1×10^5^ cells were then seeded and transduced with lentiviruses of non-silencing (NS) shRNA, single shRNAs of Cx43, PIK3CA, PIK3CB, or PIK3CD, or combinations of Cx43 shRNA and one PI3K shRNA. Cells were selected with 0.5 μg/mL puromycin for 72 hours, with media changes each day. Cells were then ready for drug treatment assays.

### Gene overexpression

pBABE-PIK3CA-E545K, pCMV5-ERK2-L73PS151D, and pBABE-SRC-Y527F were purchased from Addgene. Transfection and expression of these plasmids were described previously (56).

### ATP/glutamate release

ATP release was measured using the Kinase-Glo^®^ Luminescent Kinase Assay (Promega) as per the manufacturer’s instructions. Glutamate release was measured using the Amplex™ Red Glutamic Acid/Glutamate Oxidase Assay Kit (ThermoFisher) according to the manufacturer’s instruction. In brief, SF-295 cells were plated at 4 x 10^4^ cells/mL in a 24-well plate and allowed to attach. Cells were then treated with vehicle (DMSO) and αCT1 (100 μM) over a 24-hour period with 100 μL of supernatant collected at 0, 6, 12, 18, and 24-hour time points in triplicate. 25 μL of Kinase-Glo^®^ reagent was mixed with the samples and incubated for 10 minutes at room temperature (RT). Luminescence was read on a FilterMax F3 microplate reader (Molecular Devices, LLC) according to the manufacturer’s instructions. Glutamate release was measured using the Amplex™ Red Glutamic Acid/Glutamate Oxidase Assay Kit (ThermoFisher). In brief, LN229 and LN229/GSC cells were plated at 1 x 10^4^ cells/mL in a 24-well plate and allowed to attach. Cells were then treated with vehicle (DMSO), TMZ (50 μM) once, and αCT1 (100 μM) every 24 hours for a total of 2 doses in triplicate. After 48 hours of treatment, 50 μL of supernatant was collected and mixed with 50 μL of Amplex^®^ Red reagent and incubated for 30 minutes at RT. The sample fluorescence was read on a FilterMax F3 microplate reader with excitation of 560 nm and emission at 590 nm. The fold change of the Kinase-Glo^®^ luminescence or the Amplex^®^ Red fluorescence was defined as the ratio of the relative Kinase-Glo^®^ luminescence or the Amplex^®^ Red fluorescence in the treated cells to that of the control cells.

### CDI calculations

CDI was calculated using the formula: Survival rate of the combination / (Survival rate of treatment 1 x Survival rate of treatment 2), based on previous reports (36–39).

### Mouse experiments

Mouse experiments were performed based on the methods described previously (30, 56, 60), with modifications. All animal studies were approved by the Institutional Animal Care and Use Committee of Virginia Tech. 2 x 10^6^ SF-295 cells were mixed with Matrigel^®^ Matrix (Corning) and subcutaneously injected into the flanks of 8-week-old SCID/beige mice (Taconic Biosciences). 8 days post injection, mice were treated with drugs as indicated in the figure. Drugs were administered every other day via intraperitoneal injection (TMZ and TGX-221) or through intratumoral injection (αCT1). Tumors were measured daily using a caliper. On day 18, mice were euthanized and tumors were harvested. Tumor volumes (mm^3^) were calculated using the formula: (length x width^2^)/2.

## Statistical analyses

One-way ANOVA, Fisher’s exact test, and student *t* test were used to determine statistical significance.

## Authors’ Contributions

Conception and design: Z.S., K.J.P., and R.G.G.

Development of methodology: K.J.P., F.S., S.G. S.L., J.J., R.V., and Z.S.

Acquisition of data (performed experiments, provided reagents, etc.): K.J.P., F.S., K.L.S., S.G., M.L., P.K., S.L., G.L, M.M., J.J., R.V., and D.F.K.

Analysis and interpretation of data (e.g., statistical analysis, computational analysis): K.J.P., K.L.S., R.V., and Z.S.

Assistance in data interpretation: C.L.G. and G.G.G.

Writing, review, and/or revision of the manuscript: Z.S., R.G.G., and K.J.P.

## Acknowledgements

This study is supported by National Institutes of Health (NIH) R21 grants R21CA216768 and R21CA245631 to Z.S., NIH R01 grants HL56728 and HL141855 to R.G.G., St Baldrick’s Foundation Summer Medical Student Fellowships to F.S. and P.K., and Translational Neurobiology Summer Undergraduate Research Fellowships from the Fralin Biomedical Research Institute to M.M and G.L. The results shown in Figure 1 are in part based upon data generated by the TCGA Research Network: https://www.cancer.gov/tcga.

## Supplemental Data

**Supplemental Table S1.**
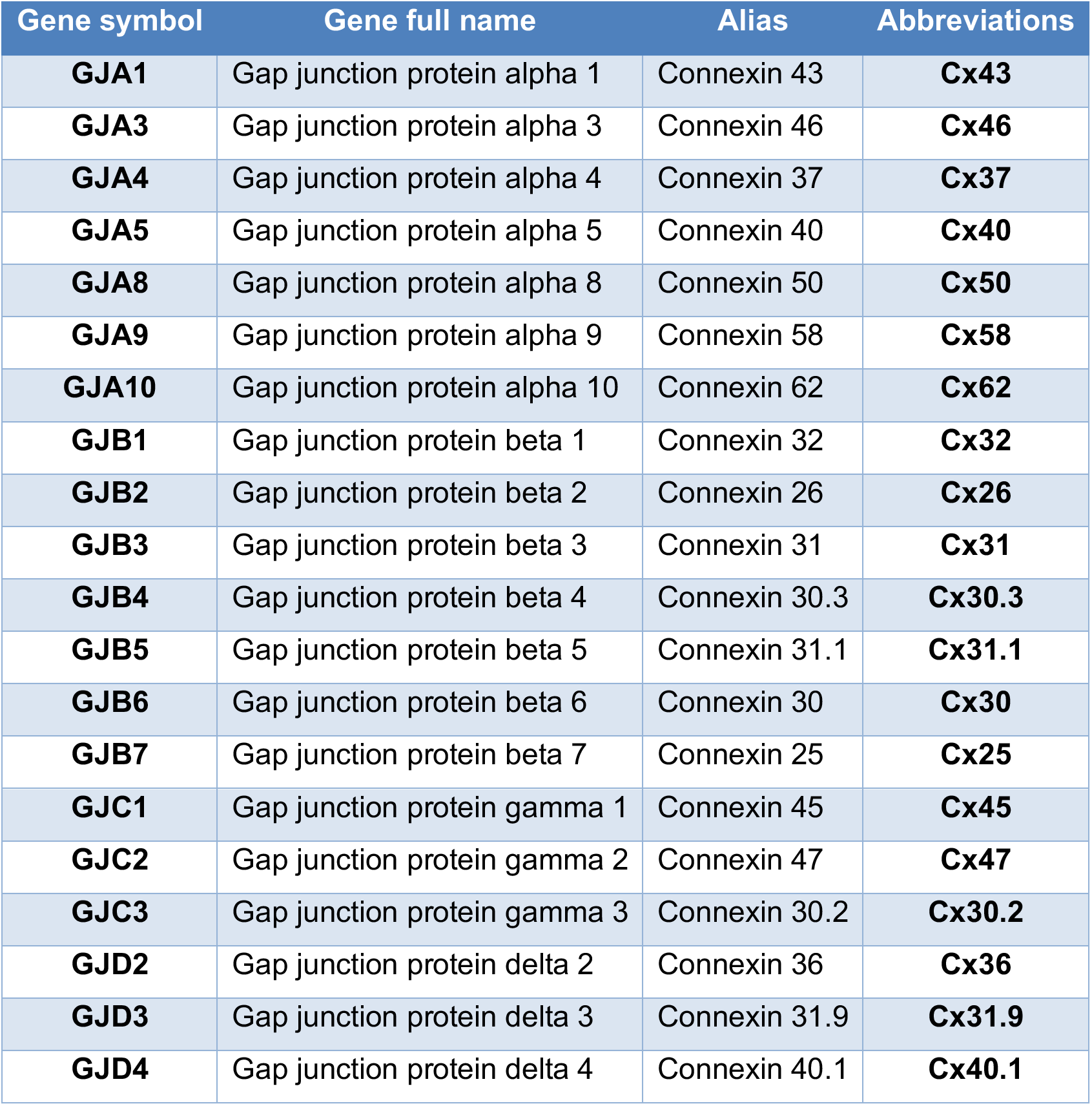
Nomenclature of connexins. Information regarding gene symbols and aliases was retrieved from GeneCards (https://www.genecards.org).

| Gene symbol | Gene full name | Alias | Abbreviations |
| --- | --- | --- | --- |
| **GJA1** | Gap junction protein alpha 1 | Connexin 43 | **Cx43** |
| **GJA3** | Gap junction protein alpha 3 | Connexin 46 | **Cx46** |
| **GJA4** | Gap junction protein alpha 4 | Connexin 37 | **Cx37** |
| **GJA5** | Gap junction protein alpha 5 | Connexin 40 | **Cx40** |
| **GJA8** | Gap junction protein alpha 8 | Connexin 50 | **Cx50** |
| **GJA9** | Gap junction protein alpha 9 | Connexin 58 | **Cx58** |
| **GJA10** | Gap junction protein alpha 10 | Connexin 62 | **Cx62** |
| **GJB1** | Gap junction protein beta 1 | Connexin 32 | **Cx32** |
| **GJB2** | Gap junction protein beta 2 | Connexin 26 | **Cx26** |
| **GJB3** | Gap junction protein beta 3 | Connexin 31 | **Cx31** |
| **GJB4** | Gap junction protein beta 4 | Connexin 30.3 | **Cx30.3** |
| **GJB5** | Gap junction protein beta 5 | Connexin 31.1 | **Cx31.1** |
| **GJB6** | Gap junction protein beta 6 | Connexin 30 | **Cx30** |
| **GJB7** | Gap junction protein beta 7 | Connexin 25 | **Cx25** |
| **GJC1** | Gap junction protein gamma 1 | Connexin 45 | **Cx45** |
| **GJC2** | Gap junction protein gamma 2 | Connexin 47 | **Cx47** |
| **GJC3** | Gap junction protein gamma 3 | Connexin 30.2 | **Cx30.2** |
| **GJD2** | Gap junction protein delta 2 | Connexin 36 | **Cx36** |
| **GJD3** | Gap junction protein delta 3 | Connexin 31.9 | **Cx31.9** |
| **GJD4** | Gap junction protein delta 4 | Connexin 40.1 | **Cx40.1** |

**Supplemental Table S2.** Levels of Cx43, pAKT-S473, p110β, MGMT and TMZ IC50. Data were retrieved from our previous publications (21, 30).

| Cell lines | Cx43 | pAKT-S473 | p110β | MGMT | TMZ IC50 (μM) |
| --- | --- | --- | --- | --- | --- |
| SF295 | 1.4 | 2.0 | 0.5 | No | 500 |
| U87MG | 3.2 | 2.2 | 0.7 | No | 1000 |
| A172 | 0.5 | 0.1 | 0.4 | No | 30 |
| LN229 | 0.5 | 0.0 | 0.2 | No | 20 |
| SF268 | 0.7 | 0.1 | 0.3 | No | 20 |
| SNB75 | 0.9 | 0.2 | 0.2 | No | 293 |

**Supplemental Fig. S1.**
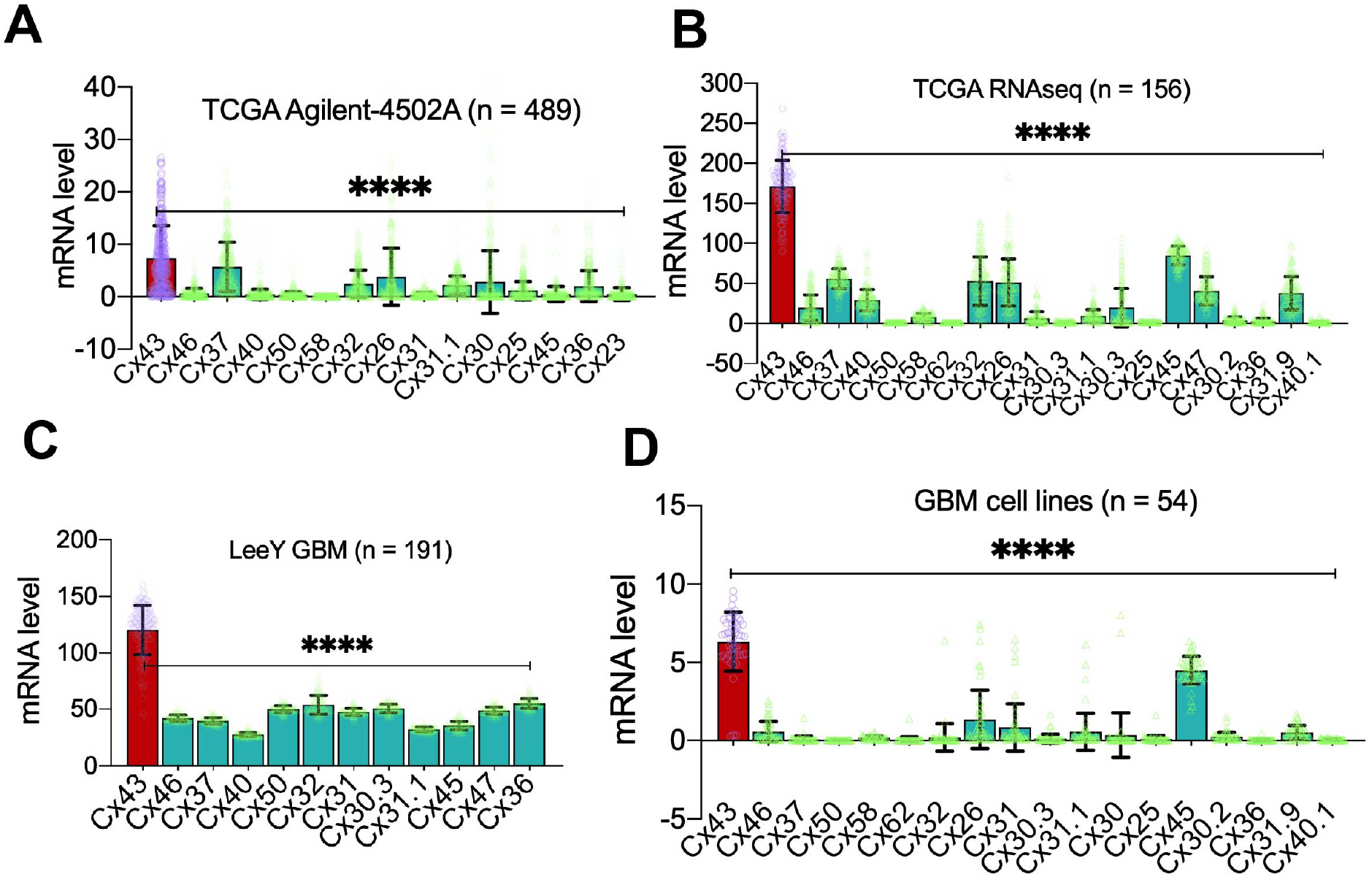
mRNA levels of connexins in GBM. Gene expression data were retrieved from cBioPortal, GlioVis, or the Cancer Dependency Map (DepMap). Shown are mRNA levels of connexins in the TCGA Agilent-4502A microarray (**A**), the TCGA RNAseq (**B**), the LeeY GBM dataset (**C**), and DepMap GBM cell lines (**D**). Case numbers (n) are also shown. Error bars represent standard deviations. Cx43 is highlighted in red and other connexins are in green. Individual data points are also shown (purple for Cx43 and yellow for other connexins). *P* values were obtained using One-Way ANOVA ****: *P* < 0.0001.

**Supplemental Fig. S2.**
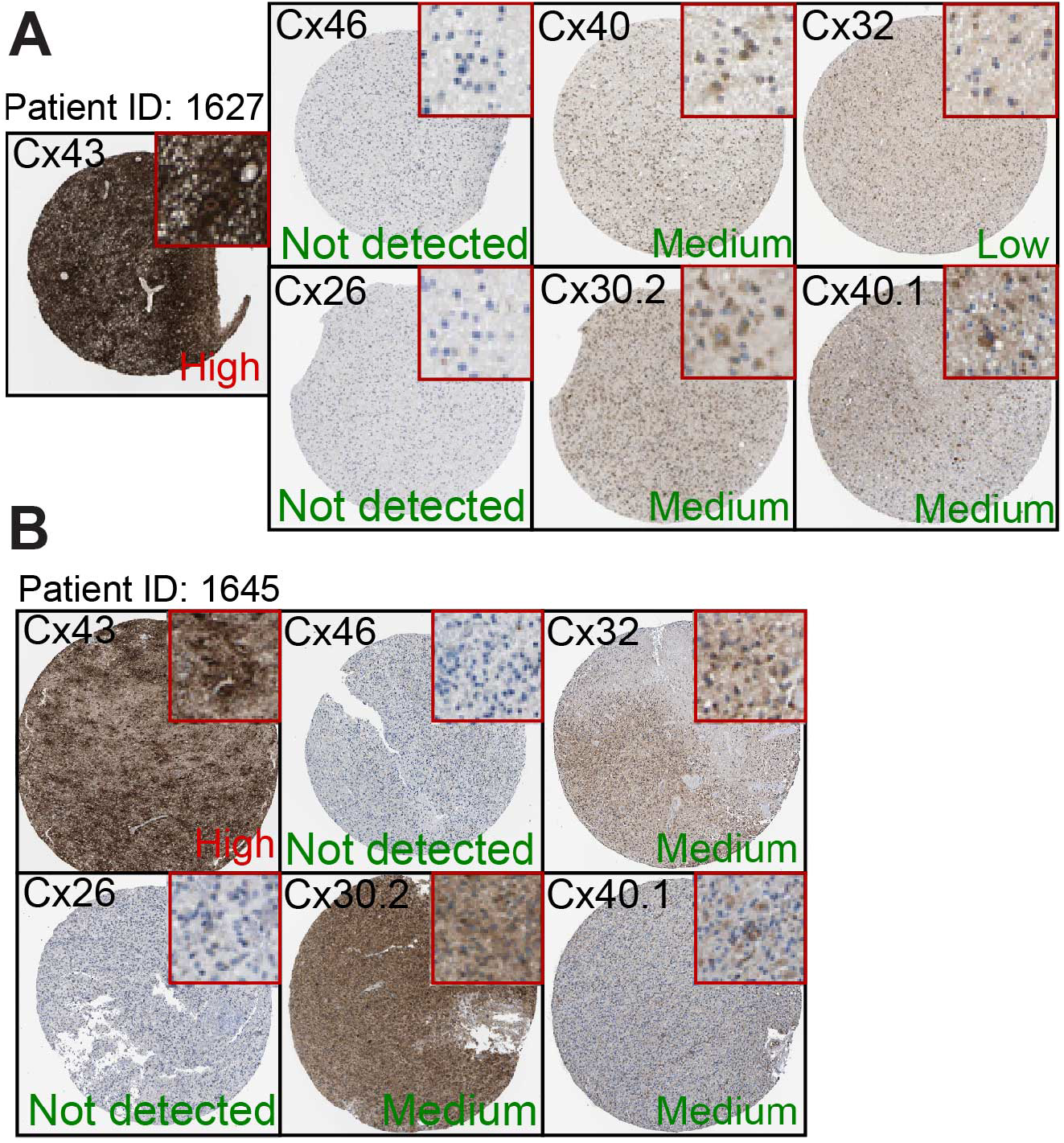
Levels of connexins in high-grade glioma. Immunohistochemical staining images of high-grade glioma were retrieved the Human Protein Atlas. Images of two patient specimens are shown in **A** and **B**, respectively. Inset figures depict details of immunostaining. Levels of staining are highlighted in red (Cx43) or in green (other connexins).

**Supplemental Fig. S3.**
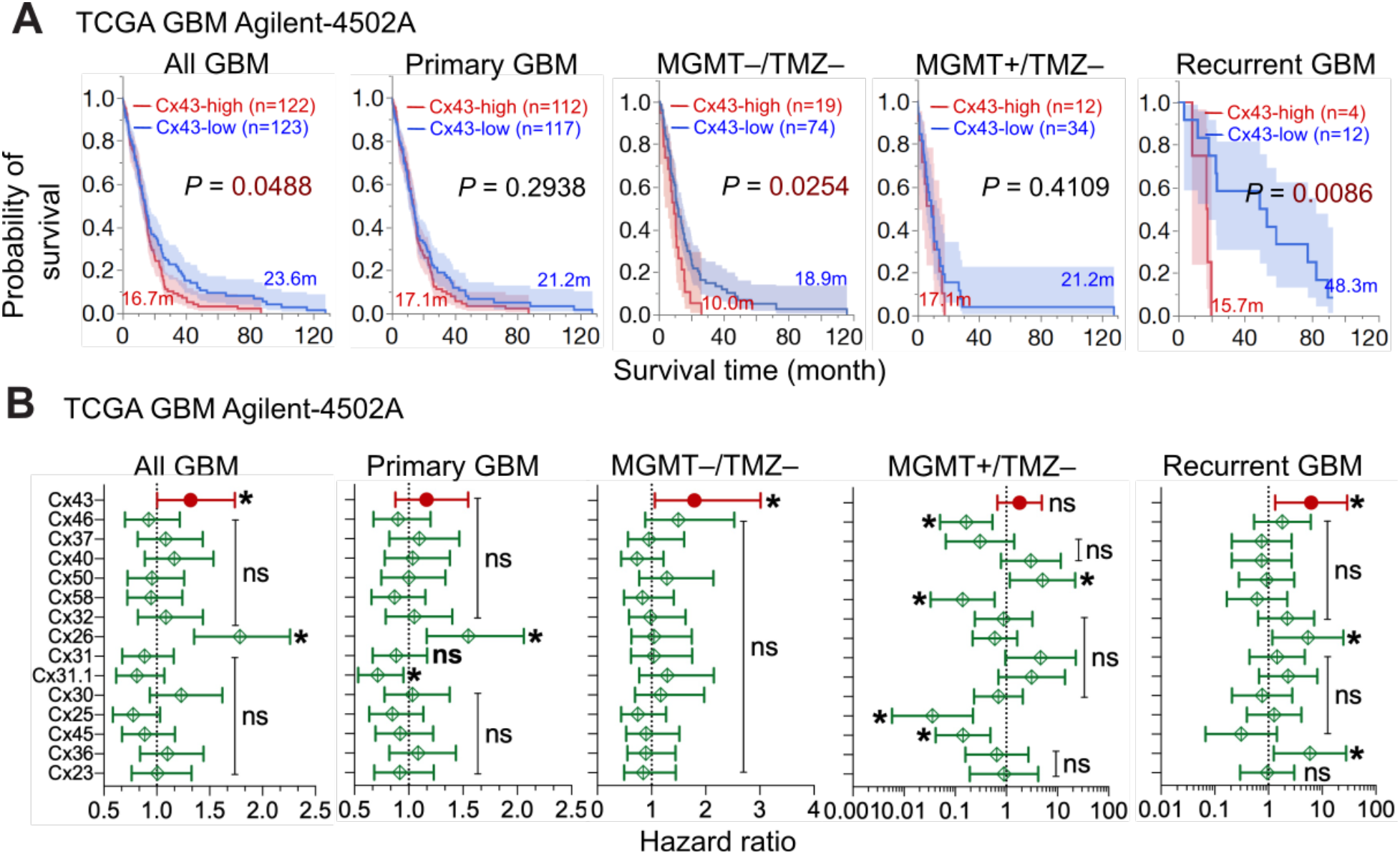
Kaplan-Meier analysis and Cox univariate analysis in the TCGA Agilent-4502A dataset. Data were retrieved from cBioportal. Patients were divided into Cx43-high (red, top 25 percentile) and Cx43-low (blue, bottom 25 or 75 percentile) based upon Cx43 mRNA levels in primary, secondary, and recurrent GBM (All GBM), primary GBM only (Primary GBM), MGMT-deficient/TMZ-untreated primary GBM (MGMT–/TMz–), MgMT-expressing/TMZ-untreated primary GBM (MGMT–/TMZ–), or recurrent GBM only (Recurrent GBM). Kaplan-Meier analysis (**A**) and Cox univariate analysis (**B**) were used. Case number (n), average survival time in months (m), 95% CI (shadow), long-rank *P* values, and hazard ratios are shown. *: *P* < 0.05. ns: not significant.

**Supplemental Fig. S4.**
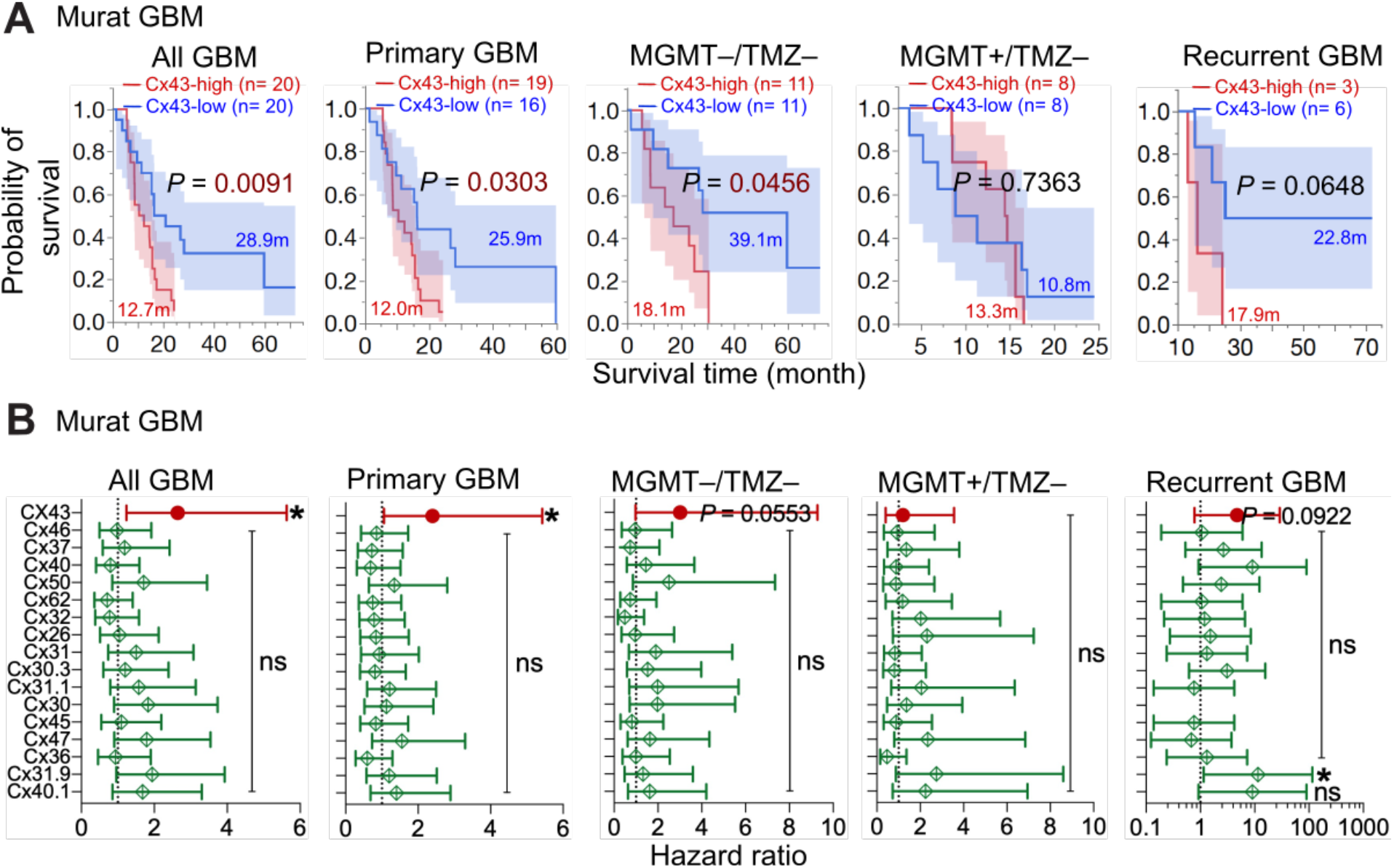
Kaplan-Meier analysis and Cox univariate analysis in the Murat GBM dataset. Data were retrieved from GlioVis. Patients were divided into Cx43-high (red, top 25 percentile) and Cx43-low (blue, bottom 25 or 75 percentile) based upon Cx43 mRNA levels in primary, secondary, and recurrent GBM (All GBM), primary GBM only (Primary GBM), MGMT-deficient/TMZ-untreated primary GBM (MGMT–/TMZ–), MGMT-expressing/TMZ-untreated primary GBM (MGMT–/TMZ–), or recurrent GBM only (Recurrent GBM).. Kaplan-Meier analysis (**A**) and Cox univariate analysis (**B**) were used. Case number (n), average survival time in months (m), 95% CI (shadow), long-rank *P* values, and hazard ratios are shown. *: *P* < 0.05. ns: not significant.

**Supplemental Fig. S5.**
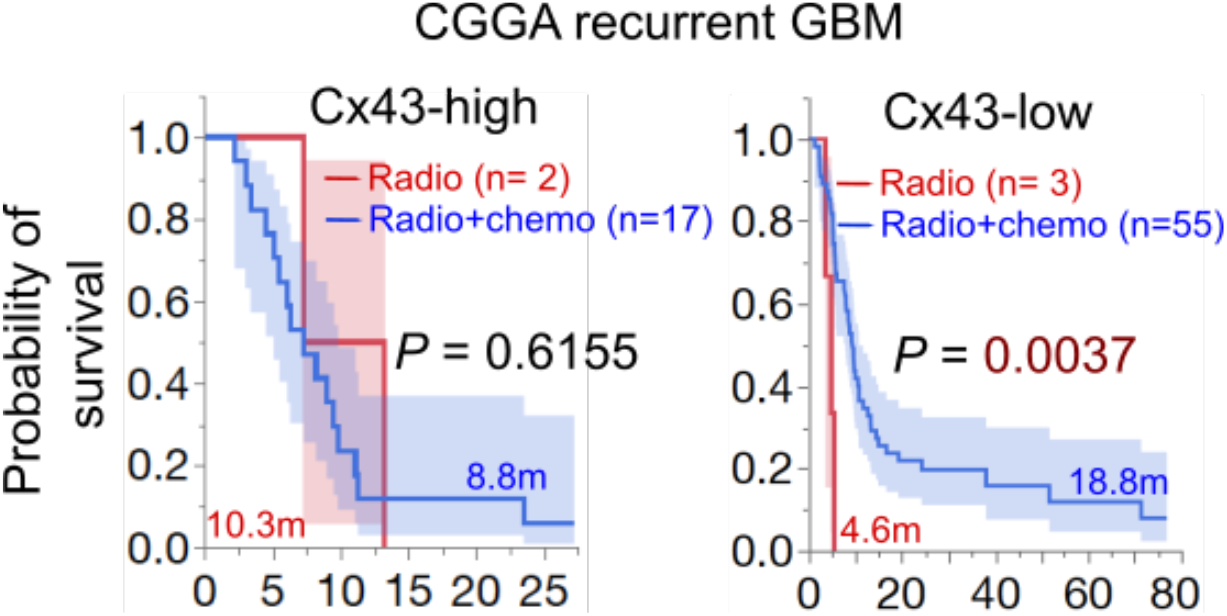
Kaplan-Meier analysis in the CGGA recurrent GBM dataset. Data were retrieved from the CGGA data protal. Cx43-hgih (top 25 percentile) or Cx43-low (bottom 75 percentile) patients were divided into Radio (red, treated with radiation only) or Radio+chemo (blue, treated with radiation and chemotherapy) based on Cx43 mRNA levels in recurrent GBMs. Case number (n), average survival time in months (m), long-rank *P* values, and hazard ratios are shown. *: *P* < 0.05 and ns: not significant.

**Supplemental Fig. S6.**
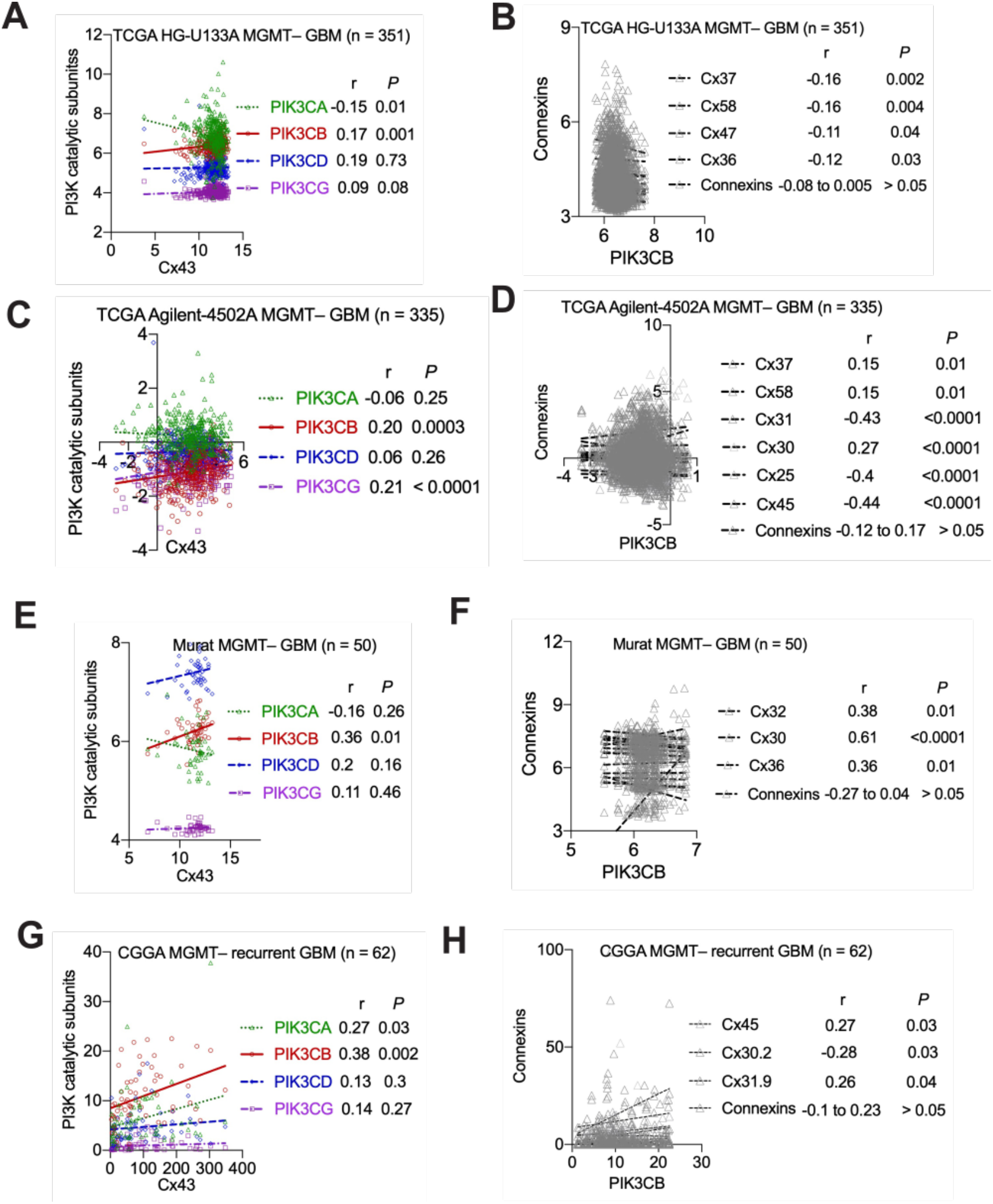
Correlation between connexins and PI3K catalytic subunits. Gene expression data were analyzed using the Pearson correlation coefficient assay. mRNA levels of Cx43 were compared to mRNA levels of PI3K catalytic subunits (**A, C, E**, and **G**) in four different datasets as indicated. mRNA levels of PIK3CB were compared to those of connexin mRNAs (**B, D, F**, and **H**). The coefficient r and corresponding *P* values are shown.

**Supplemental Fig. S7.**
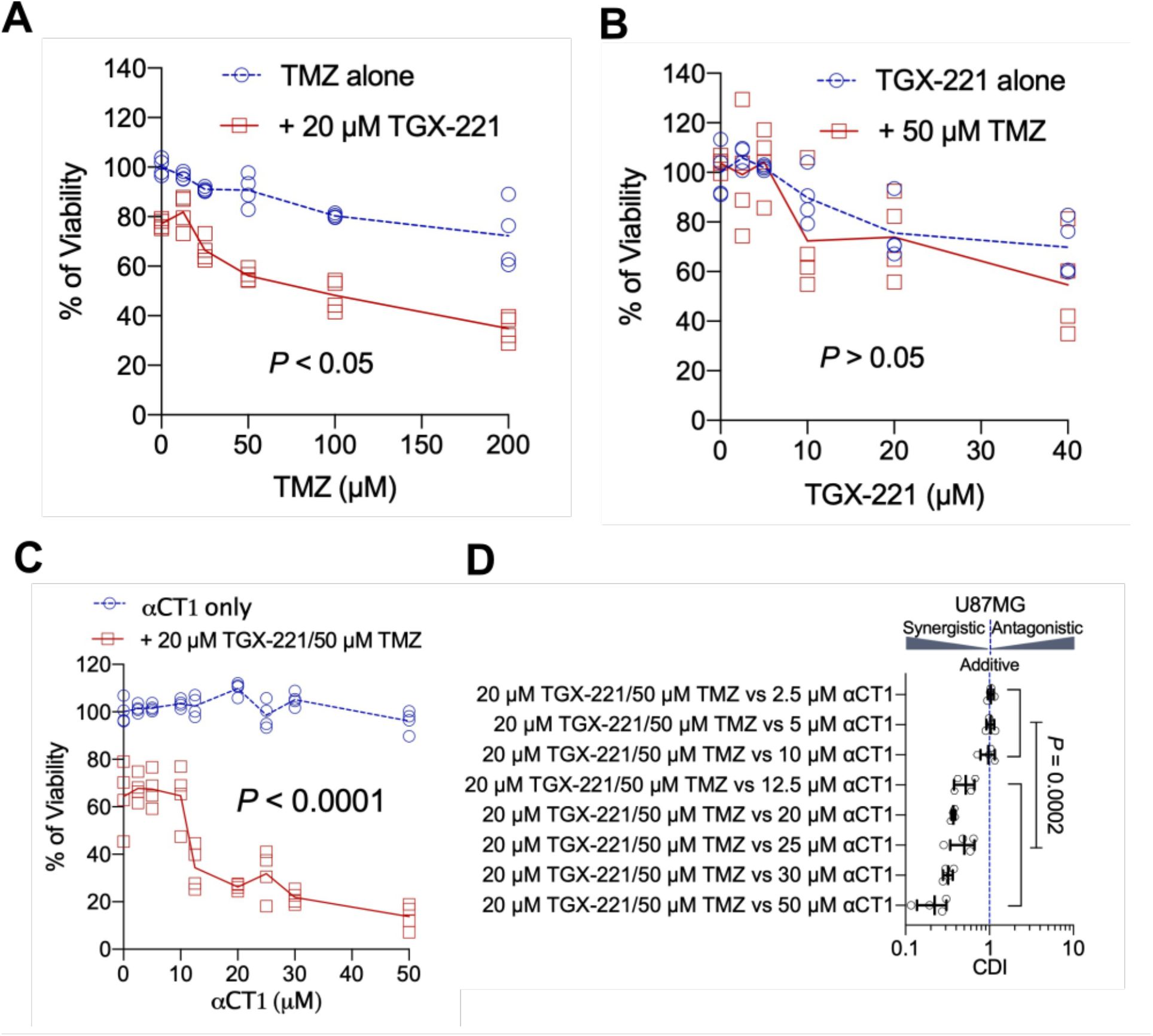
Optimization of αCT1, TGX-221 and TMZ in U87MG cells. (**A**) Combination of 20 μM TGX-221 and TMZ at various concentrations. U87MG cells were treated with drug combinations as indicated for 6 days. Cell viability was determined using the MTS viability assay. The vehicle DMSO was the control and set as 100%. Treated cells were normalized to DMSO-treated cells. (**B**) Combination of 50 μM TMZ and TGX-221 at various concentrations. (**C**) Combination of 20 μM TGX-221/50 μM TMZ and αCT1 at different concentrations. (**D**) CDIs of different combinations tested in **C**. One-way ANOVA and student *t* test were used to determine statistical significance.

**Supplemental Fig. S8.**
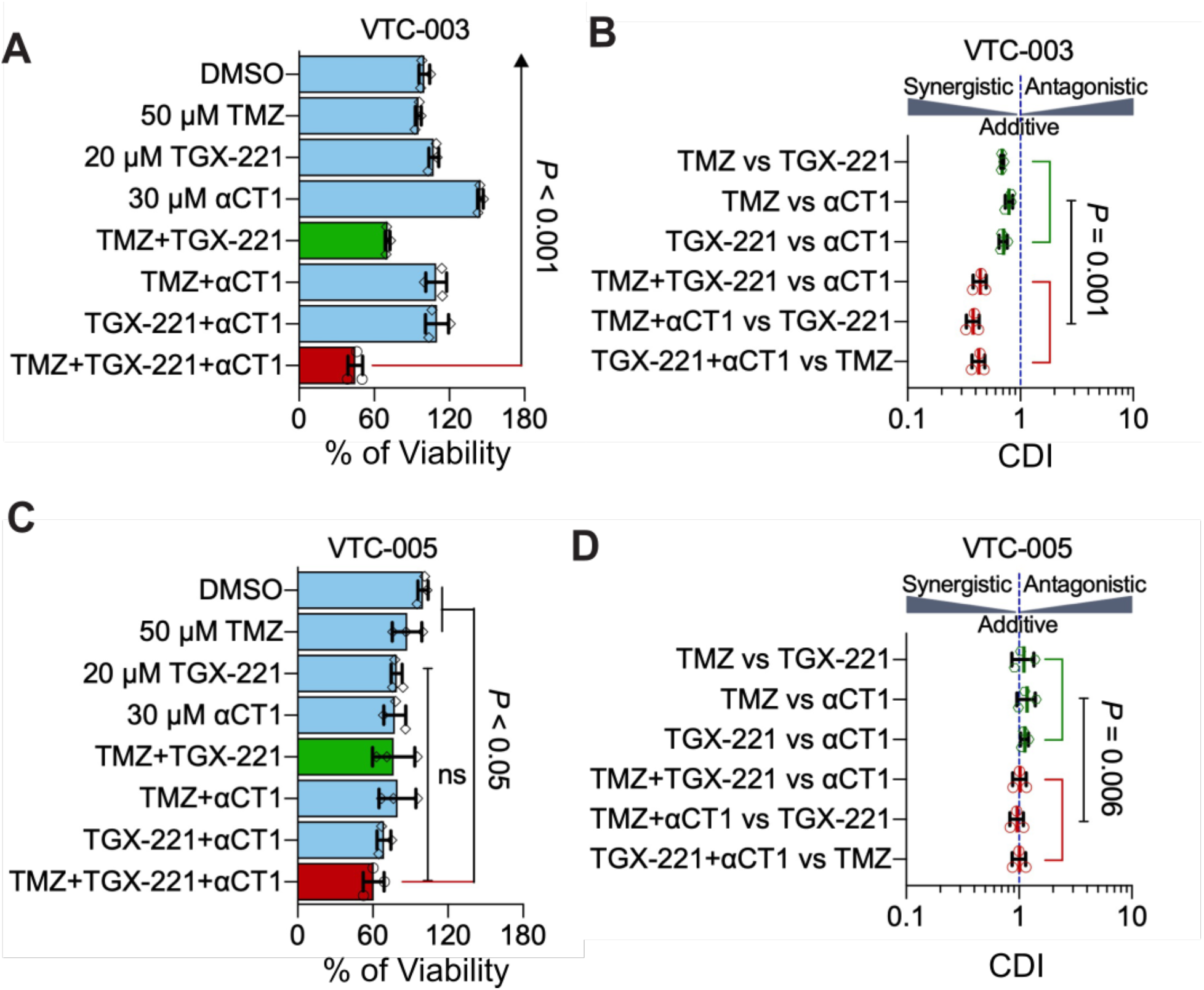
The αCT1/TGX combo in VTC-003 and VTC-005. (**A**) Viability of VTC-003 cells treated with different drug combinations for 6 days. Cell viability was determined using the MTS viability assay. (**B**) CDIs of drug combinations tested in VTC-003 cells. (**C**) Viability of VTC-005 cells treated with different drug combinations. (**D**) CDIs of drug combinations tested in VTC-005. One-way ANOVA or student *t* test were used to determine statistical significance.

**Supplemental Fig. S9.**
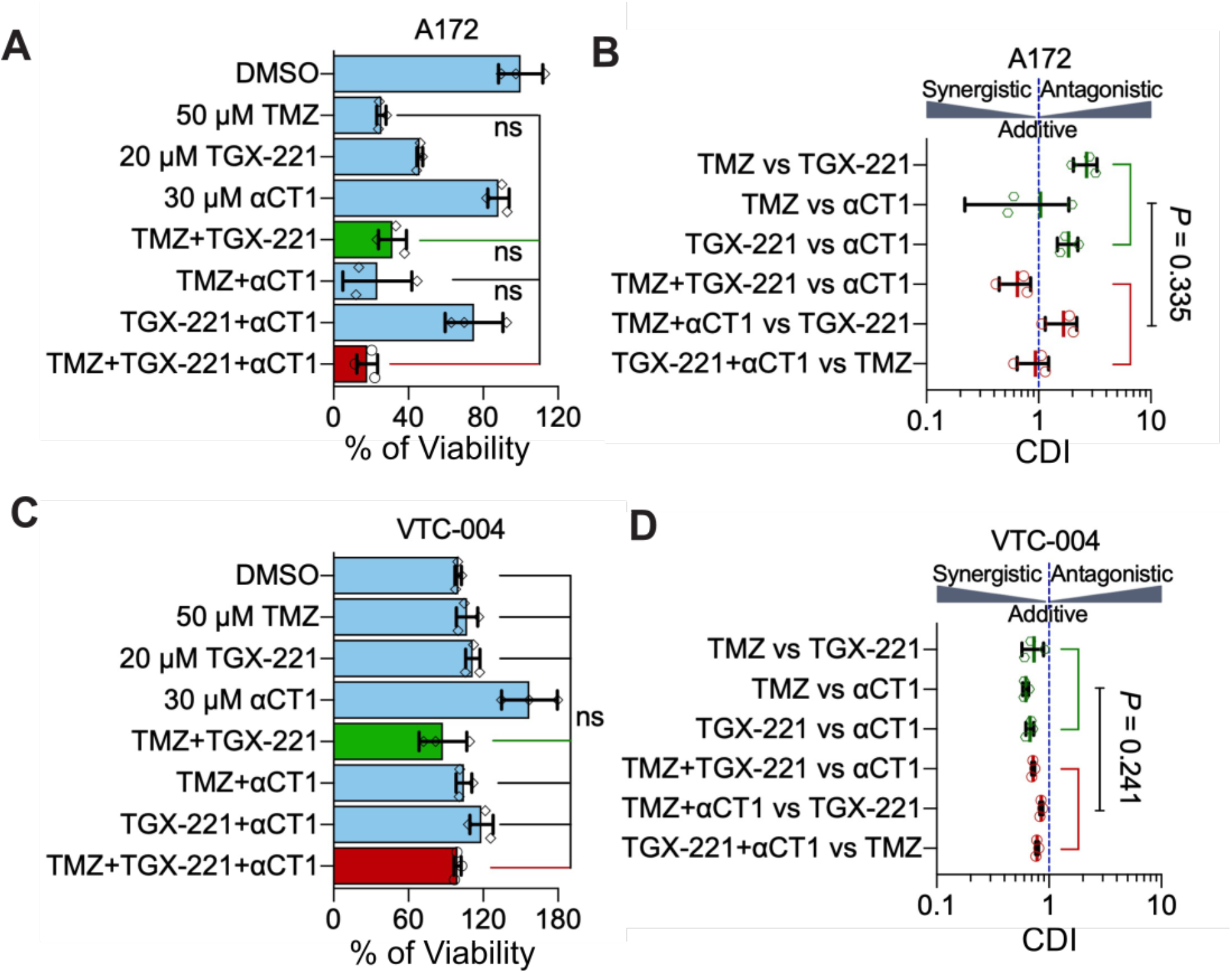
The αCT1/TGX combo in A172 and VTC-004. (**A**) Viability of A172 cells treated with different drug combinations for 6 days. Cell viability was determined using the MTS viability assay. (**B**) CDIs of drug combinations tested in A172. (**C**) Viability of VTC-004 cells treated with different drug combinations. (**D**) CDIs of drug combinations tested in VTC-004. One-way ANOVA or student *t* test were used to determine statistical significance.

**Supplemental Fig. S10.**
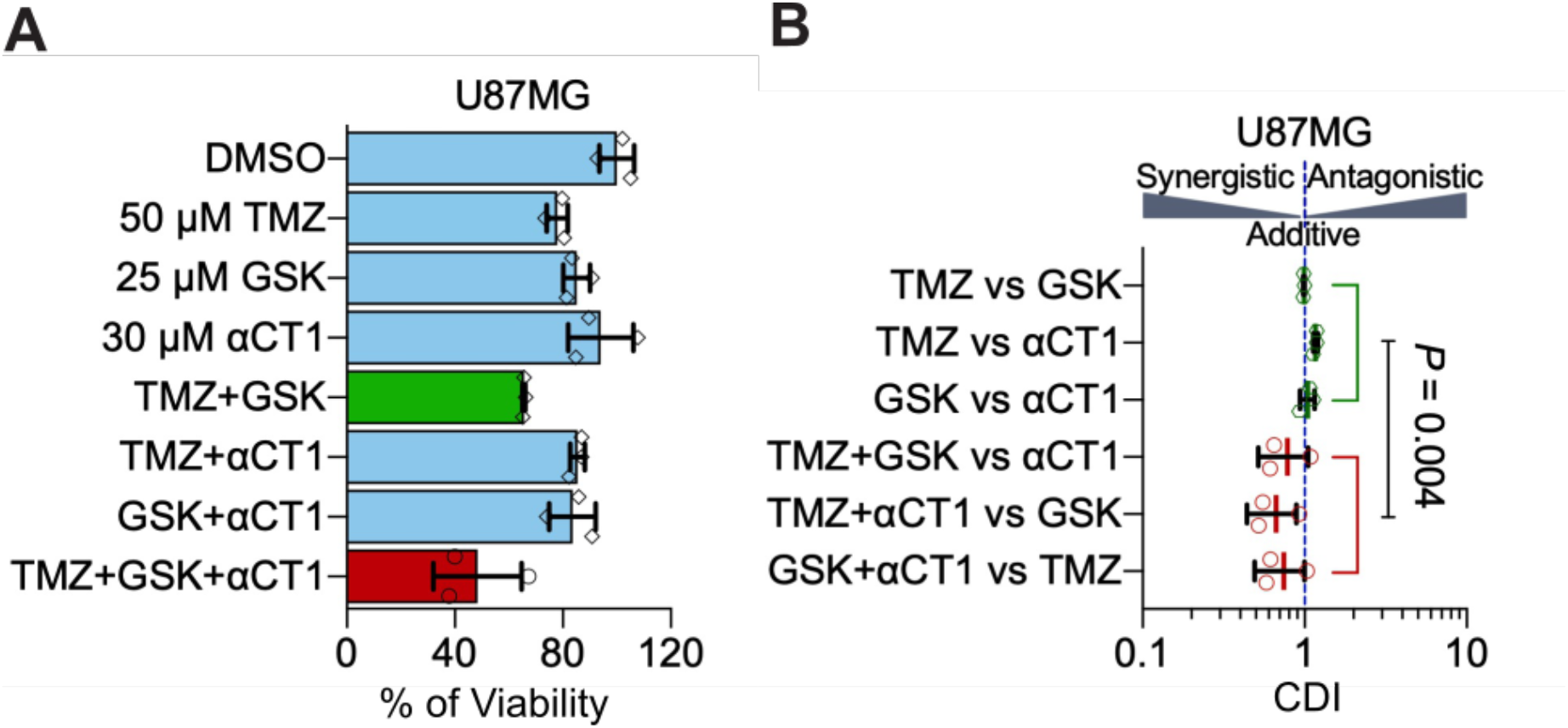
The αCT1/GSK combo in U87MG cells. (**A**) Viability of U87MG cells treated with different combinations of drugs. GSK: GSK2636771 for 6 days. Cell viability was determined using the MTS viability assay. (**B**) CDIs of drug combinations tested in U87MG. Student *t* test were used to determine statistical significance.

